# Diffuse optical reconstructions of fNIRS data using Maximum Entropy on the Mean

**DOI:** 10.1101/2021.02.22.432263

**Authors:** Zhengchen Cai, Alexis Machado, Rasheda Arman Chowdhury, Amanda Spilkin, Thomas Vincent, Ümit Aydin, Giovanni Pellegrino, Jean-Marc Lina, Christophe Grova

## Abstract

Functional near-infrared spectroscopy (fNIRS) measures the hemoglobin concentration changes associated with neuronal activity. Diffuse optical tomography (DOT) consists of reconstructing the optical density changes measured from scalp channels to the oxy-/deoxy-hemoglobin (i.e., HbO/HbR) concentration changes within the cortical regions. In the present study, we adapted a nonlinear source localization method developed and validated in the context of Electro- and Magneto-Encephalography (EEG/MEG): the Maximum Entropy on the Mean (MEM), to solve the inverse problem of DOT reconstruction. We first introduced depth weighting strategy within the MEM framework for DOT reconstruction to avoid biasing the reconstruction results of DOT towards superficial regions. We also proposed a new initialization of the MEM model improving the temporal accuracy of the original MEM framework. To evaluate MEM performance and compare with widely used depth weighted Minimum Norm Estimate (MNE) inverse solution, we applied a realistic simulation scheme which contained 4000 simulations generated by 250 different seeds at different locations and 4 spatial extents ranging from 3 to 40*cm*^2^ along the cortical surface. Our results showed that overall MEM provided more accurate DOT reconstructions than MNE. Moreover, we found that MEM was remained particularly robust in low signal-to-noise ratio (SNR) conditions. The proposed method was further illustrated by comparing to functional Magnetic Resonance Imaging (fMRI) activation maps, on real data involving finger tapping tasks with two different montages. The results showed that MEM provided more accurate HbO and HbR reconstructions in spatial agreement with the main fMRI cluster, when compared to MNE.

**Highlights:**

- We introduced a new fNIRS reconstruction method - Maximum Entropy on the Mean.
- We implemented depth weighting strategy within the MEM framework.
- We improved the temporal accuracy of the original MEM reconstruction.
- Performances of MEM and MNE were evaluated with realistic simulations and real data.
- MEM provided more accurate and robust reconstructions than MNE.

## 1. Introduction

Functional Near-infrared spectroscopy (fNIRS) is an non-invasive functional neuroimaging modality. It detects changes in oxy-/deoxy-hemoglobin (i.e., HbO/HbR) concentration within head tissues through the measurement of near-infrared light absorption using sources and detectors placed on the surface of the head (Scholkmann et al., 2014; Yücel et al., 2021). In continuous wave fNIRS, the conventional way to transform variations in optical density to HbO/HbR concentration changes at the level of each source-detector channel, is to apply the modified Beer Lambert Law (mBLL) (Delpy et al., 1988). This model assumes homogeneous concentration changes within the detecting region, i.e., ignoring the partial volume effects which indicates the absorption of light within the illuminated regions varies locally. This assumption reduces quantitative accuracy of HbO/HbR concentration changes when dealing with focal hemodynamic changes (Boas et al., 2001; Strangman et al., 2003).

In order to handle these important quantification biases associated with sensor level based analysis, diffuse optical tomography (DOT) has been proposed to reconstruct, from sensor level measures of the optical density, the fluctuations of HbO/HbR concentrations within the brain (Arridge, 1999). This technique not only provides better spatial localization accuracy and resolution of the underlying hemodynamic responses (Boas et al., 2004a; Joseph et al., 2006), but also avoids partial volume effect in classical mBLL, hence achieves better quantitative estimation of HbO/HbR concentration changes (Boas et al., 2001; Strangman et al., 2003). DOT has been applied to reconstruct hemodynamic responses in sensory and motor cortex during median-nerve stimulation (Dehghani et al., 2009; Hughes et al., 2004) and finger tapping (Boas et al., 2004a; Yamashita et al., 2016); to conduct visual cortex retinotopic mapping (Zeff et al., 2007; White and Culver, 2010; Eggebrecht et al., 2012) and to simultaneous image hemodynamic responses over the motor and visual cortex (White et al., 2009).

To formalize DOT reconstruction, one needs to solve two main problems. The first one is the forward problem which estimates a forward model or sensitivity matrix that maps local absorption changes within the brain to variations of optical density changes measured by each channel (Boas et al., 2002). The second problem is the inverse problem which aims at reconstructing the fluctuations of hemodynamic activity within the brain from scalp measurements (Arridge, 2011). The forward problem can be solved by generating a subject specific anatomical model, describing accurately propagation of light within the head. Such anatomical model is obtained by segmenting anatomical Magnetic Resonance Imaging (MRI) data, typically into five tissues (i.e., scalp, skull, cerebro-spinal fluid (CSF), white matter and gray matter), before initializing absorption and scattering coefficients values for each tissue type and for each wavelength (Fang, 2010; Machado et al., 2018). Solving the inverse problem relies on solving an ill-posed problem which does not provide a unique solution, unless specific additional constraints are added. The most widely used inverse method in DOT is a linear approach based on Minimum Norm Estimate (MNE) originally proposed for solving the inverse problem of MagnetoencephaloGraphy(MEG) and Electroencephalography (EEG) source localization (Hämäläinen and Ilmoniemi, 1994). It minimizes the *L*_2_ norm of the reconstruction error along with Tikhonov regularization (Boas et al., 2004b; Zeff et al., 2007; Dehghani et al., 2009; Eggebrecht et al., 2012, 2014; Tremblay et al., 2018). Other strategies to solve DOT inverse problem have also been considered, such as sparse regularization using the *L*_1_ norm (Süzen et al., 2010; Okawa et al., 2011; Kavuri et al., 2012; Prakash et al., 2014; Tremblay et al., 2018) and Expectation Maximization (EM) algorithm (Cao et al., 2007). A non-linear method based on hierarchical Bayesian model for which inference is obtained through an iterative process (Shimokawa et al., 2012, 2013) has been proposed and applied on finger tapping experiments in (Yamashita et al., 2016).

Maximum Entropy on the Mean (MEM) framework was first proposed by Amblard et al., 2004 and then applied and carefully evaluated by our group in the context of EEG/MEG source imaging (Grova et al., 2006; Chowdhury et al., 2013). The MEM framework was specifically designed and evaluated for its ability to recover spatially extended generators (Heers et al., 2016; Pellegrino et al., 2016; Chowdhury et al., 2016; Grova et al., 2016). We recently demonstrated its excellent performances when dealing with focal sources (Hedrich et al., 2017) and when applied on clinical epilepsy data (Chowdhury et al., 2018; Pellegrino et al., 2020). In addition to its unique ability to recover the spatial extent of the underlying generators, we also demonstrated MEM’s excellent accuracy in low SNR conditions, with the ability to limit the influence of distant spurious sources (Chowdhury et al., 2016; Hedrich et al., 2017; Heers et al., 2016; Pellegrino et al., 2020; von Ellenrieder et al., 2016; Aydin et al., 2020).

We believe that these important aspects should be carefully considered in the context of fNIRS reconstruction. The first one is the ability to accurately recover the spatial extent of the underlying hemodynamic activity for both focal and extended generators. The second one is to provide robust reconstruction results when data SNR decreases, especially when considering the fact that it is challenging to maintain a good intra-subject consistence using continuous-wave fNIRS due to its relatively low SNR (Chen et al., 2020). Therefore, our main objective was to adapt the MEM framework for fNIRS reconstruction and carefully evaluate its performance. Moreover, fNIRS reconstruction results tends to be biased towards more superficial regions, because the light sensitivity profile decreases exponentially with the depth of the generators (Strangman et al., 2013). To overcome this bias, we implemented and evaluated a depth weighted variant of the MEM framework.

The article is organized as follows. The methodology of depth weighted MEM for DOT is first presented. Then, we described our validation framework using realistic simulations and associated validation metrics. fNIRS reconstruction using MEM was compared with widely used depth weighted Minimum Norm Estimate (MNE) inverse solution. Finally, illustrations of the methods on finger tapping fNIRS data set acquired with two different montages from 6 healthy subjects are provided and compared with functional Magnetic Resonance Imaging (fMRI) results.

## 2. Material and Methods

### 2.1. fNIRS reconstruction

To perform fNIRS reconstructions, the relationship between measured optical density changes on the scalp and wavelength specific absorption changes within head tissue is usually expressed using the following linear model (Arridge, 1999):

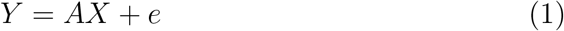

where *Y* is a matrix (*p* × *t*) which represents the wavelength specific measurement of optical density changes in *p* channels at *t* time samples. *X* (*q* × *t*) represents the unknown wavelength specific absorption changes in *q* locations along the cortex at time *t*. *A* (*p* × *q*) is called the light sensitivity matrix which is actually the forward model relating absorption changes in the head to optical density changes measured in each channel. Finally, *e* (*p* × *t*) models the additive measurement noise. Solving the fNIRS tomographic reconstruction problem consists in solving an inverse problem which can be seen as the estimation of matrix *X* (i.e. the amplitude for each location *q* at time *t*). However, this problem is ill-posed and admits an infinite number of possible solutions. Therefore, solving the DOT inverse problem requires adding additional prior information or regularization constraints to identify a unique solution.

In DOT studies, anatomical constraints can be considered by defining the reconstruction solution space (i.e. where *q* is located) within the gray matter volume (Boas and Dale, 2005) or along the cortical surface (Huppert et al., 2017; Machado et al., 2021). In EEG and MEG source localization studies (Dale and Sereno, 1993; Grova et al., 2006; Chowdhury et al., 2013), it also is common to constrain the reconstruction along the cortical surface. In this study, the reconstruction space was considered as the mid surface defined as the middle layer between gray matter/pial and gray/white matter interfaces (Fischl et al., 2002).

### 2.2. Minimum Norm Estimation (MNE)

Minimum norm estimation is one of the most widely used reconstruction methods in DOT (Zeff et al., 2007; Dehghani et al., 2009; White et al., 2009; White and Culver, 2010; Eggebrecht et al., 2012, 2014; Yamashita et al., 2016). Such estimation can be expressed using a Bayesian formulation which solves the inverse problem by estimating the posterior distribution 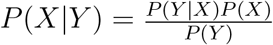(i.e. the probability distribution of parameter *X* conditioned on data *Y*). A solution can be computed by imposing Gaussian distribution priors on the generators 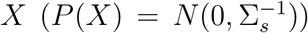 and the noise 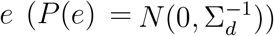. Σ_*d*_ is the inverse of the noise covariance which could be estimated from baseline recordings. Σ_*s*_ is the inverse of the source covariance which is assumed to be an identity matrix in conventional MNE.

The Maximum a Posteriori (MAP) estimator of the posterior distribution *P*(*X|Y*) can be obtained using maximum likelihood estimation:

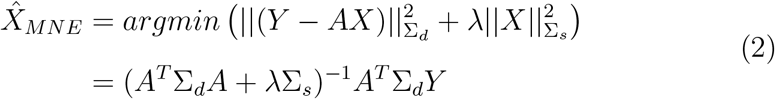

where 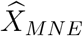 is the reconstructed absorption changes along the cortical surface. λ is a hyperparameter to regularize the inversion using the priori minimum norm constraint 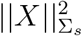. In this study, we applied the standard L-Curve method to estimate this λ as suggested in (Hansen, 2000).

### 2.3. Depth weighted MNE

Standard MNE solutions assumes Σ_*s*_ = *I*, which then tends to bias the inverse solution towards the generators exhibiting large sensitivity in the forward model, therefore the most superficial ones (Fuchs et al., 1999). When compared to EEG-MEG source localization, such bias is even more pro-nounced in fNIRS since within the forward model light sensitivity values decrease exponentially with the depth (Strangman et al., 2013). This bias can be compensated by scaling the source covariance matrix such that the variances are equalized (van der Sluis, 1969; Fuchs et al., 1999). In the context of DOT, depth weighted MNE has been proposed by Culver et al., 2003 as an approach to compensate this effect and applied in different studies (Zeff et al., 2007; Dehghani et al., 2009; White et al., 2009; Eggebrecht et al., 2012, 2014). In practice, depth weighting can be formulated differently, here we consider a generalized expression for the implementation of depth weighted MNE as proposed in Lin et al., 2006. It consists in initializing the source covariance matrix as 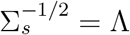, resulting in a so called depth weighted MNE solution, described as follows:

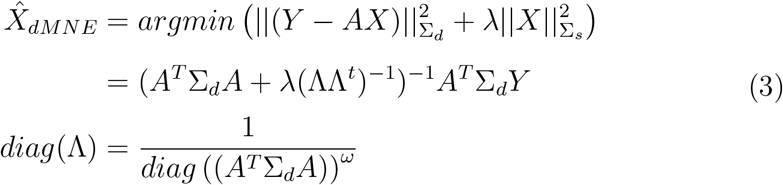

Depth weighted MNE solution takes into account the forward model *A* for each position in the brain and therefore penalizes most superficial regions exhibiting larger amplitude in *A*, by enhancing the contribution to deeper regions. *ω* is a weighting parameter tuning the amount of depth compensation to be applied. The larger is *ω*, the more depth compensation is considered. *ω* = 0 would therefore refer to no depth compensation and an identity source covariance model. *ω* = 0.5 refers to standard depth weighting approach mentioned above. In the present study, we carefully evaluated the impact of this parameter on DOT accuracy with a set of *ω* values (i.e. *ω* = 0,0.1,0.3, 0.5, 0.7 *and* 0.9).

### 2.4. Maximum Entropy on the Mean (MEM) for fNIRS 3D reconstruction

#### 2.4.1. MEM framework

The main contribution of this study is the first adaptation and evaluation of MEM method (Amblard et al., 2004; Grova et al., 2006; Chowdhury et al., 2013) to perform DOT reconstructions in fNIRS. Within the MEM framework, the intensity of *x*, i.e. amplitude of *X* at each location *q* in Eq. 1, is considered as a random variable, described by the following probability distribution *dp*(*x*) = *p*(*x*)*dx*. The Kullback-Leibler divergence or *ν*-entropy of *dp*(*x*) relative to a prior distribution *dν*(*x*) is defined as,

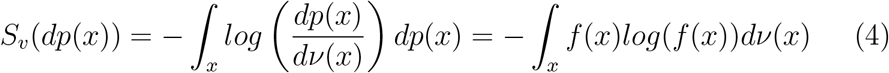

where *f*(*x*) is the *ν*-density of *dp*(*x*) defined as *dp*(*x*) = *f*(*x*)*dν*(*x*). Following a Bayesian approach to introduce the data fit, we denote *C_m_* as the set of probability distributions on *x* that explains the data on average:

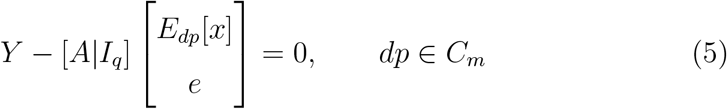

where *Y* represents the measured optical density changes, 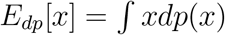 represents the statistical expectation of *x* under the probability distribution *dp*, and *I_q_* is an identity matrix of (*q* × *q*) dimension. Therefore, within the MEM framework, a unique solution of *dp*(*x*) could be obtained,

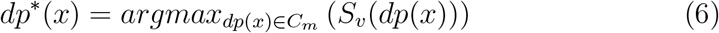

The solution of *dp*\*(*x*) can be solved by maximizing the *ν*-entropy which is a convex function. It is equivalent to minimize an unconstrained concave Lagrangian function i.e., *L*(*dp*(*x*), *κ*, λ), along with two Lagrangian constraint parameters, i.e., *κ* and λ. It is finally equivalent to maximize a cost function *D*(λ) which is described as,

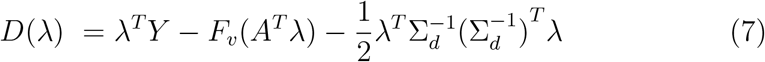

where 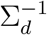 is the noise covariance matrix. *F_υ_* represents the free energy associated with reference *dν*(*x*). It is important to mention that *D*(λ) is now an optimization problem within a space of dimension equal to the number of sensors. Therefore, if we estimate λ* = *argmax*_λ_*D*(λ), the unique solution of MEM framework is then obtained from the gradient of the free energy.

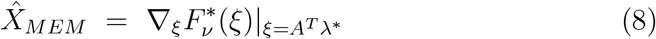

For further details on MEM implementation and theory we refer the reader to (Amblard et al., 2004; Grova et al., 2006; Chowdhury et al., 2013).

#### 2.4.2. Construction of the prior distribution for MEM estimation

To define the prior distribution *dν*(*x*) mentioned above, we assumed that brain activity can be depicted by a set of K non-overlapping and independent cortical parcels. Then the reference distribution *dν*(*x*) can be modeled as,

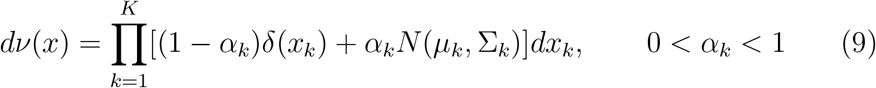

Each cortical parcel k is characterized by an activation state, defined by the hidden variable *S_k_*, describing if the parcel is active or not. Therefore we denote *α_k_* as the probability of *k^th^* parcel to be active, i.e., *Prob*(*S_k_* = 1). *δ_k_* is a Dirac function that allows to “switch off” the parcel when considered as inactive (i.e., *S_k_* = 0). *N*(*μ_k_*, Σ_*k*_) is a Gaussian distribution, describing the distribution of absorptions changes within the *k^th^* parcel, when the parcel is considered as active (*S_k_* = 1). This prior model, which is specific to our MEM inference, offers a unique opportunity to switch off some parcels of the model, resulting in accurate spatial reconstructions of the underlying activity patterns with their spatial extent, as carefully studied and compared with other Bayesian methods in Chowdhury et al., 2013.

The spatial clustering of the cortical surface into K non-overlapping parcel was obtained using a data driven parcellization (DDP) technique (Lapalme et al., 2006). DDP consisted in first applying a projection method, the multivariate source prelocalization (MSP) technique (Mattout et al., 2005), estimating a probability like coefficient (MSP score) between 0 and 1 for each vertex of the cortical mesh, characterizing its contribution to the data. DDP is then obtained by using a region growing algorithm, along the tessellated cortical surface, starting from local MSP maxima. Once the parcellization is done, the prior distrubution *dν*(*x*) is then a joint distribution expressed as the multiplication of individual distribution of each parcel in Eq. 9 assuming statistical independence between parcels,

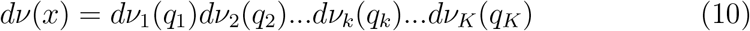

where *dν*(*x*) is the joint probability distribution of the prior, *dν_k_*(*q_k_*) is the individual distribution of the parcel *k* described as Eq. 9.

To initialize the prior in Eq. 9, *μ_k_* which is the mean of the Gaussian distribution, *N*(*μ_k_*, Σ_*k*_), was set to zero. Σ_*k*_ at each time point *t*, i.e. Σ_*k*_(*t*), was defined by Eq. 11 according to (Chowdhury et al., 2013),

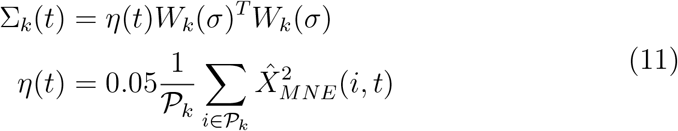

where *W_k_*(*σ*) is a spatial smoothness matrix, defined by (Friston et al., 2008), which controls the local spatial smoothness within the parcel according to the geodesic surface neighborhood order. Same value of *σ* = 0.6 was used as in (Chowdhury et al., 2013). *η*(*t*) was defined as 5% of the averaged energy of MNE solution within each parcel 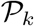 at time *t*. Finally, we can substitute this initialization into Eq. 9 to construct the prior distribution *dν*(*x*), and then obtain the MEM solution using Eq. 8.

It is worth mentioning that we did not use MNE solution as the prior of *μ_k_* in Eq. 9 at all, which was actually initialized to 0 in our framework. We only used 5% of the averaged energy of MNE solution, over the parcel *k*, to set the prior for covariance Σ_*k*_. The posterior estimation of parameter *μ_k_* was estimated from the Bayesian framework by conditioning with data. Moreover, the prior of MEM framework is a mixture of activation probability *α_k_* and a Gaussian distribution (see Eq. 9), in which the prior for *α_k_* was informed by a spatio-temporal extension of the MSP score (see Chowdhury et al., 2013 for further details). These aspects completely differentiate MEM from approaches that iteratively update reconstruction results initialized by a MNE solution.

#### 2.4.3. Depth weighted MEM

In addition to adapting MEM for fNIRS reconstruction, we also implemented for the first time, depth weighting within the MEM framework. Two depth weighting parameters, *ω*_1_ and *ω*_2_, were involved in this process. *ω*_1_ was used to apply depth weighting on the source covariance matrix Σ_*k*_ of each parcel *k* in Eq. 11. *ω*_2_ was applied to solve the depth weighted MNE, as described in Eq. 3, before using those prior to initialize the source covariance model within each parcel of the MEM model. Therefore, the standard MNE solution 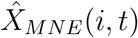 in Eq. 11 was replaced by the depth weighted version of MNE solution 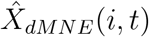 described by Eq. 3. Consequently, the depth weighted version of Σ_*k*_(*t*) is now defined as,

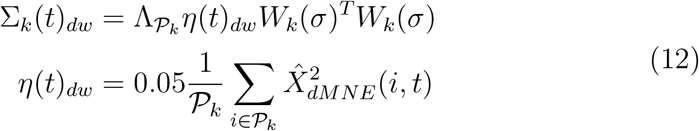

where 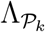 is the depth weighting matrix for each pacel *k*, in which *ω*_1_ was involved to construct this scaling matrix as described in Eq. 3. This initialization followed the logic that depth weighting is in fact achieved by scaling the source covariance matrix. The other depth weighting parameter, *ω*_2_, was considered when solving 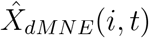, therefore avoiding biasing the initialization of the source covariance with a standard MNE solution.

To comprehensively compare MEM and MNE and also to investigate the behavior of depth weighting, we first evaluated the reconstruction performance of MNE with different *ω*_2_ (i.e. step of 0.1 from 0 to 0.9). Then two of these values (i.e. *ω*_2_ = 0.3 and 0.5) were selected for the comparison with MEM since they performed better than the others. Note that the following expressions of depth weighted MEM will be denoted as MEM(*ω*_1_, *ω*_2_) to represent the different depth weighting strategies.

#### 2.4.4. Accuracy of temporal dynamics

The last contribution of this study was to improve the temporal accuracy of MEM solutions. In classical MEM approach (Chowdhury et al., 2013), 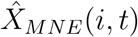 in Eq. 12 was globally normalized by dividing by max 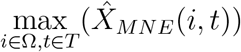 where Ω represents all the possible locations along the cortical surface and *T* is the whole time segment. Therefore, the constructed prior along the time actually contained the temporal scaled dynamics from MNE solution. To remove this effect, we performed local normalization for 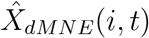 at each time instance *t*, i.e., by dividing by 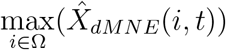. This new feature would preserve the spatial information provided by prior distribution, while allowing MEM to estimate the temporal dynamics only from the data.

### 2.5. Validation of fNIRS reconstruction methods

We evaluated the performance of the two fNIRS reconstruction methods (i.e., MEM and MNE), first within a fully controlled environment involving the use of realistic simulations of fNIRS data, followed by evaluations on real data acquired with a well controlled finger tapping paradigm. Two different fNIRS montages were considered in those two proposed evaluations.

**Montage 1**: A full Double Density (DD) montage (see Fig. 1) which is a widely used fNIRS montage, was considered given that it allows sufficient dense spatial coverage of fNIRS channel to allow local DOT (Kawaguchi et al., 2007). One healthy subject underwent fNIRS acquisitions with this DD montage, involving the two following sessions,

- A 10 minutes resting state session was acquired to add realistic physiology noise to be considered in our realistic simulations. The subject was seating on a comfortable armchair and instructed to keep the eyes open and to remain awake. The optodes of the full DD montage (i.e. 8 sources and 10 detectors resulting in 50 fNIRS channels) are presented in Fig. 1e. The montage composed of 6 second-order distance channels (1.5*cm*), 24 third-order channels(3*cm*) and 12 fourth-order channels with 3.35*cm* distance. In addition, we also added one proximity detector paired for each source to construct close distance channels (0.7*cm*) in order to measure superficial signals within extra-cerebral tissues. To place the montage with respect to the region of interest, the center of the montage was aligned with the center of the right “hand knob” area, which controls the left hand movement (Raffin et al., 2015), projected on the scalp surface and then each optodes were projected on the scalp surface (see Fig. 1d).
- The subject was asked to sequentially tap the left thumb against the other digits around 2Hz, therefore the main elicited hemodynamic response was indeed expected over the right hand knob area. The finger tapping paradigm consisted in 10 blocks of 30s tapping task and each of them was followed by a 30 to 35s resting period. The beginning/end of each block was informed by an auditory cue.
**Montage 2**: A personalized optimal montage (see Fig. 8) following the methodology we previously reported in Machado et al., 2018. First, the hand knob within right primary motor cortex was drawn manually along the cortical surface and defined as a target region of interest (ROI) using the Brainstorm software (Tadel et al., 2011). Then we applied optimal montage estimation (Machado et al., 2014, 2018) in order to estimate personalized montages, built to maximize a priori fNIRS sensitivity and spatial overlap between channels with respect to the target ROI. To ensure good spatial overlap between channels for local 3D reconstruction, we constructed personalized optimal montages composed of 3 sources and 15 detectors (see Fig. 7b). The source-detector distance was set to vary from 2*cm* to 4.5*cm* and each source was constrained such that it has to create channels with at least 13 detectors. Finally, we also manually added 1 proximity channel, located at the center of the 3 sources. Five subjects underwent fNIRS acquisitions with personalized optimal montage during a similar finger tapping task as the one for montage 1, in which 20 blocks were acquired by alternating a task (period of 10*s*) and a resting state period ranging from 30*s* to 60*s*.

**Fig. 1.**
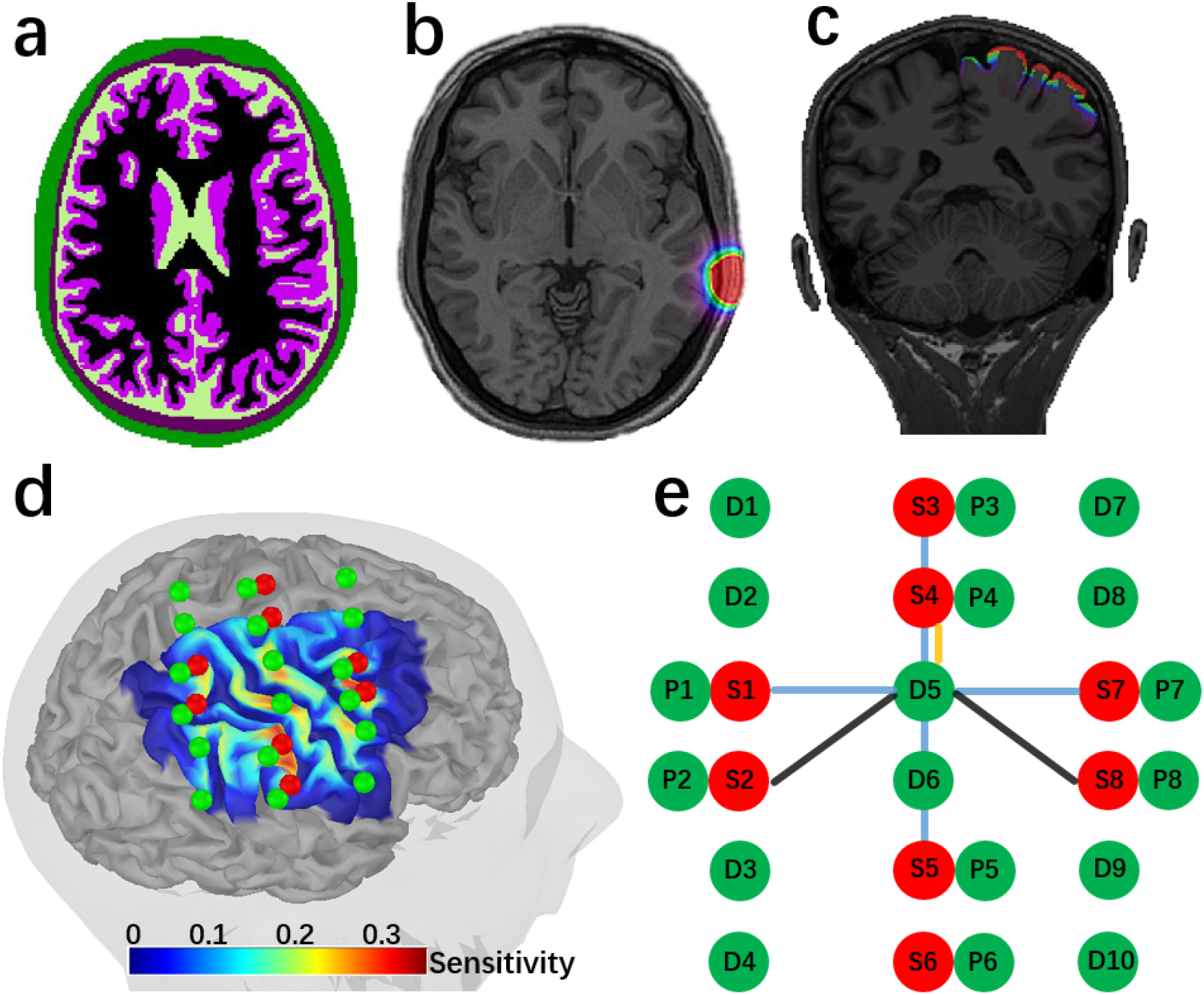
fNIRS measurement montage 1 and the anatomical model considered for DOT forward model estimation. (a) Anatomical 3D MRI segmented in five tissues, namely, scalp (green), skull (brown), CSF (light green), gray matter (purple) and white matter (black). (b) Optical fluence of one optode calculated through Monte Carlo simulation of Photons within this head model, using MCXLab. (c) Sensitivity profile of the whole montage in volume space. (d) Sensitivity profile, i.e. the summation of sensitivity map of all channels, along the cortical surface. Green dots represent detectors, including one proximity detector 0.7 cm for each source, and red dots represent sources. (e) double-density montage 1 considered for this acquisition. There were 50 channels in total, 12 of 3.8cm (black), 24 of 3cm (blue), 6 of 1.5cm (yellow) and 8 of close distance (0.7*cm*) channels.

All 6 subjects have signed written informed consent forms for this study which was approved by the Central Committee of Research Ethics of the Minister of Health and Social Services Research Ethics Board, Québec, Canada.

#### 2.5.1. MRI and fMRI Data acquisitions

Anatomical MRI data were acquired on those 6 healthy subjects (25 ± 6 years old, right-handed) and were considered to generate realistic anatomical head models. MRI data were acquired in a GE 3T scanner at the PERFORM Center of Concordia University, Montréal, Canada. T1-weighted anatomical images were acquired using the 3D BRAVO sequence (1 × 1 × 1 mm^3^, 192 axial slices, 256 × 256 matrix), whereas T2-weighted anatomical images were acquired using the 3D Cube T2 sequence (1 × 1 × 1 mm^3^ voxels, 168 sagittal slices, 256 × 256 matrix).

Participants also underwent functional MRI acquisition (without fNIRS) while performing the same finger opposition tasks considered in fNIRS. fMRI acquisition consisted in a gradient echo EPI sequence (3.7 × 3.7 × 3.7 mm^3^ voxels, 32 axial slices, TE = 25 ms, TR = 2, 000 ms). fMRI Z-maps were generated by standard first-level fMRI analysis using FEAT from FSL v6.0.0 software (Smith et al., 2004; Jenkinson et al., 2012).

#### 2.5.2. fNIRS Data acquisition

fNIRS acquisitions were conducted at the PERFORM Center of Concordia University using a Brainsight fNIRS device (Rogue Research Inc., Montréeal, Canada), equipped with 16 dual wavelength sources (685*nm* and 830*nm*), 32 detectors and 16 proximity detectors (for short distance channels). All montages (i.e., double density and optimal montages) were installed to cover the right motor cortex. Knowing a priori the exact positions of fNIRS channels estimated on the anatomical MRI of each participant, we then used a 3D neuronavigation system (Brainsight TMS navigation system, Rogue Research Inc.) to guide the installation of the sensors on the scalp. Finally, every sensor was glued on the scalp using a clinical adhesive, collodion, to prevent motion and ensure good contact to the scalp (Yücel et al., 2014; Machado et al., 2018).

#### 2.5.3. fNIRS forward model estimation

T1 and T2 weighted anatomical images were processed using FreeSurfer V6.0 (Fischl et al., 2002) and Brain Extraction Tool2 (BET2) (Smith et al., 2004) in FMRIB Software Library (FSL) to segment the head into 5 tissues (i.e. scalp, skull, Cerebrospinal fluid (CSF), gray matter and white matter see Fig. 1a).

Same optical coefficients used in (Yücel et al., 2014; Machado et al., 2018) for the two wavelengths considered during our fNIRS acquisition, 685*nm* and 830*nm*, were assigned to each tissue type mentioned above. Fluences of light for each optode (see Fig. 1b) was estimated by Monte Carlo simulations with 10^8^ photons using MCXLAB developed by Fang and Boas, 2009; Yu et al., 2018 (http://mcx.sourceforge.net/cgi-bin/index.cgi). Sensitivity values were then computed using the adjoint formulation and were normalized by the Rytov approximation (Arridge, 1999).

For each source-detector pair of our montages, the corresponding light sensitivity map was first estimated in a volume space, and then further constrained to the 3D mask of gray matter tissue (see Fig. 1c), as suggested in Boas and Dale, 2005. Then, these sensitivity values within the gray matter volume were projected along the cortical surface (see Fig. 1d and Fig. 7c) using the Voronoi based method proposed by (Grova et al., 2006). We considered the mid-surface from FreeSurfer as the cortical surface. This surface was downsampled to 25, 000 vertices. This volume to surface interpolation method has the ability to preserve sulco-gyral morphology (Grova et al., 2006). After the interpolation, the sensitivity value of each vertex of the surface mesh represents the mean sensitivity of the corresponding volumetric Voronoi cell (i.e., a set of voxels that have closest distances to a certain vertex than to all other vertices).

#### 2.5.4. fNIRS data preprocessing

Using the coefficient of variation of the fNIRS data, channels exhibiting a standard deviation larger than 8% of the signal mean were rejected (Schmitz et al., 2005; Schneider et al., 2011; Eggebrecht et al., 2012; Piper et al., 2014). Superficial physiological fluctuations were regressed out at each channel using the average of all proximity channels’ (0.7*cm*) signals (Zeff et al., 2007). All channels were then band-pass filtered between 0.01Hz and 0.1Hz using a 3rd order Butterworth filter. Changes in optical density (i.e., Δ*OD*) were calculated using the conversion to log-ratio. Finally, Δ*OD* of finger tapping data were block averaged around the task onsets. Note that since sensors were glued with collodion, we observed very minimal motion during the acquisitions. Real background signal considered to generate realistic simulations also underwent the same preprocessing.

#### 2.5.5. Realistic Simulations of fNIRS Data

We first considered realistic simulations of fNIRS data to evaluate DOT methods within a fully controlled environment. To do so, theoretical task-induced HbO/HbR concentration changes were simulated within cortical surface regions with a variety of locations, areas and depths. Corresponding optical density changes in the channel space were then computed by applying the corresponding fNIRS forward model, before adding real resting state fNIRS baseline signal as realistic physiological noise at different signal to noise ratio (SNR) levels.

As presented in Fig. 2a, we defined three sets of evenly distributed seeds within the field of view of DOT reconstruction. The locations were selected with respect to the depth relative to the skull, namely we simulated 100 “superficial seeds”, 100 “middle seeds” and 50 “deep seeds”. The cortical regions in which we simulated an hemodynamic response were generated by region growing around those seeds, along the cortical surface. To simulate generators with different spatial extents (denoted here as Se), we considered four levels of neighborhood orders, growing geodesically along the cortical surface, resulting in spatial extents ranging from Se = 3, 5, 7, 9 (corresponding areas of 3 to 40 cm^2^). For simplification, these cortical regions within which an hemodynamic response was simulated will be denoted as “generators” in this paper. For each vertex within a “generator”, a canonical Hemodynamic Response Function (HRF) was convoluted with a simulated experimental paradigm which consisted in one block of 20s task surrounded by 60s pre-/post-baseline period (Fig. 2b). Simulated HbO/HbR fluctuations within the theoretical generator (Fig. 2c) were then converted to the corresponding absorption changes of two wavelengths (i.e., 685*nm* and 830*nm*). After applying the forward model matrix A in Eq. 1, we estimated the simulated, noise free, task induced Δ*OD* in all channels.

**Fig. 2.**
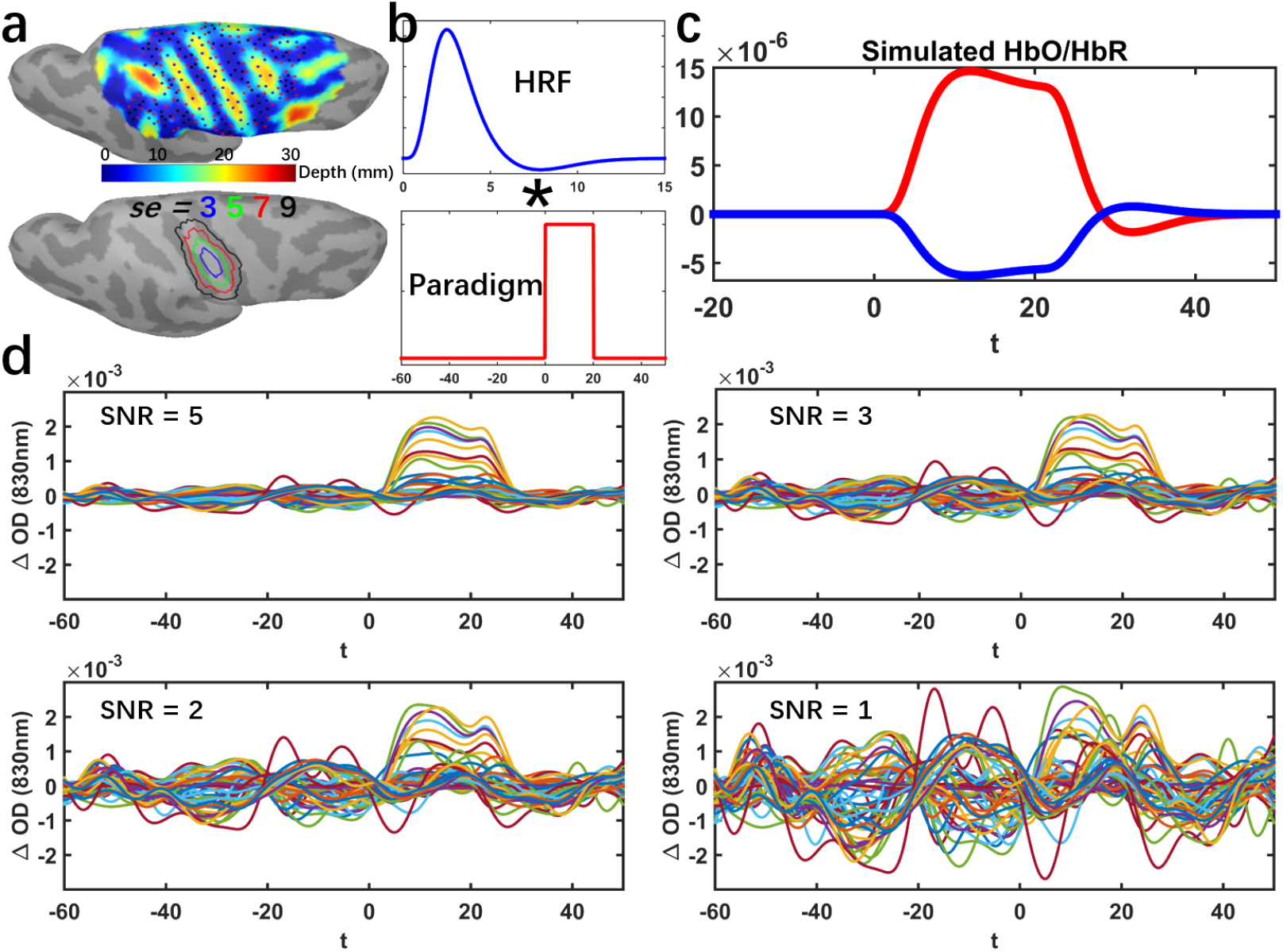
Workflow describing our proposed realistic fNIRS simulation framework.(a) 100 Superficial seeds (black dots), 100 Middle seeds (red dots), 50 Deep seeds (blue dots) with spatial extent of *Se* = 3, 5, 7, 9 neighbourhood order within the field of view. (b) Convolution of a canonical HRF model with an experimental block paradigm (60*s* before and 50*s* after the onset). (c) Simulated theoretical HbO/HbR fluctuations along the cortical surface within the corresponding generator. (d) Realistic simulations obtained by applying the fNIRS forward model and addition of the average of 10 trials of real fNIRS background measurements at 830nm. Time course of Δ*OD* of all channels with SNR of 5, 3, 2 and 1 respectively are presented

Δ*OD* of real resting state data were then used to add realistic fluctuations (noise) to these simulated signals. Over the 10min of recording, we randomly selected 10 baseline epochs of 120*s* each, free from any motion artifact by visual inspection. To mimic a standard fNIRS block average response, realistic simulations were obtained by adding the average of these 10 real baseline epochs to the theoretical noise-free simulated Δ*OD*, at five SNR levels (i.e. SNR = 5, 3, 2,1). SNR was calculated through the following equation,

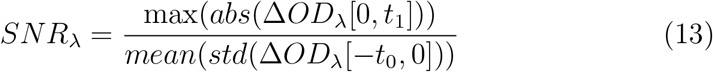

where Δ*OD*_λ_[0, *t*_1_] is the optical density changes of a certain wavelength λ in all channels during the period from 0*s* to *t*_1_ = 60*s*. *std*(Δ*OD*_λ_[−*t*_0_, 0]) is the standard deviation of Δ*OD*_λ_ during baseline period along all channels. Simulated trials for each of four different SNR levels are illustrated in Fig. 2d. A total number of 4000 realistic simulations were considered for this evaluation study, i.e., 250 (seeds) × 4 (spatial extents) × 4 (SNR levels). Note that resting state fNIRS baseline signal was preprocessed before adding to the simulated signals.

#### 2.5.6. Validation metric

Following the validation metrics described in (Grova et al., 2006; Chowdhury et al., 2013, 2016; Hedrich et al., 2017), we applied 4 quantitative metrics to access the spatial and temporal accuracy of fNIRS 3D reconstructions. Further details on the computation of those four validation metrics are reported in Supplementary material S1.

- **Area Under the Receiver Operating Characteristic (ROC) curve (AUC)** was used to assess general reconstruction accuracy considering both sensitivity and specificity. AUC score was estimated as the area under the ROC curve, which was obtained by plotting sensitivity as a function of (1-specificity). AUC ranges from 0 to 1, the higher it is the more accurate the reconstruction is.
- **Minimum geodesic distance (Dmin)** measuring the geodesic dis-tance in millimeters, following the circumvolutions of the cortical surface, from the vertex that exhibited maximum of reconstructed activity to the border of the ground truth. Low Dmin values indicate better accuracy in estimating the location of the generator.
- **Spatial Dispersion (SD)** assessed the spatial spread of the estimated generator distribution and the localization error. It is expressed in millimeters. A reconstructed map with either large spatial spread around the ground truth or large localization error would result in large SD values.
- **Shape error(SE)** evaluated the temporal accuracy of the reconstruc-tion. It was calculated as the root mean square of the difference between the normalized reconstructed time course and the normalized ground truth time course. Low SE values indicate high temporal accuracy of the reconstruction.

### 2.6. Statistics

Throughout all of the quantitative evaluations among different methods involving different depth weighting factors *ω* in the results section, Wilcoxon signed rank test was applied to test the significance of the paired differences between each comparison. For each statistical test, we reported the median value of paired differences, together with its p-value (Bonferroni corrected).

We are only showing results at 830nm for simulations, since the ones from 690nm under the same SNR level would have provided similar reconstructed spatiotemporal maps except for the reversed amplitudes. However, reconstruction results on real data indeed involved both wavelengths.

## 3. Results

### 3.1. Evaluation of MEM v.s. MNE using realistic simulations

We first investigated the effects of depth weighting factor *ω*_2_ selection for depth weighted MNE. To do so, we evaluated spatial and temporal per-formances of DOT reconstruction for a set of *ω*_2_ (step of 0.1 from 0 to 0.9). Based on those results reported in the Supplementary material S2 and Fig. S1, we decided to considered that most accurate fNIRS reconstructions were ob-tained when considering *ω*_2_ = 0.3 and 0.5 for depth weighted MNE. Therefore only those two values were further considered for comparison with MEM reconstructions.

Comparison of the performance of MEM and MNE on superficial realistic simulations are presented in Table. 1 and Fig. 3, for 4 levels of spatial extent (*Se* = 3, 5, 7, 9), using boxplot distribution of the 4 validation metrics. We evaluated 3 depth weighted implementations of MEM, namely, MEM(*ω*_1_ = 0.3, *ω*_2_ = 0.3), MEM(0.3, 0.5) and MEM(0.5,0.5), as well as 2 depth weighted implementations of MNE, namely, MNE(0.3) and MNE(0.5).

**Fig. 3.**
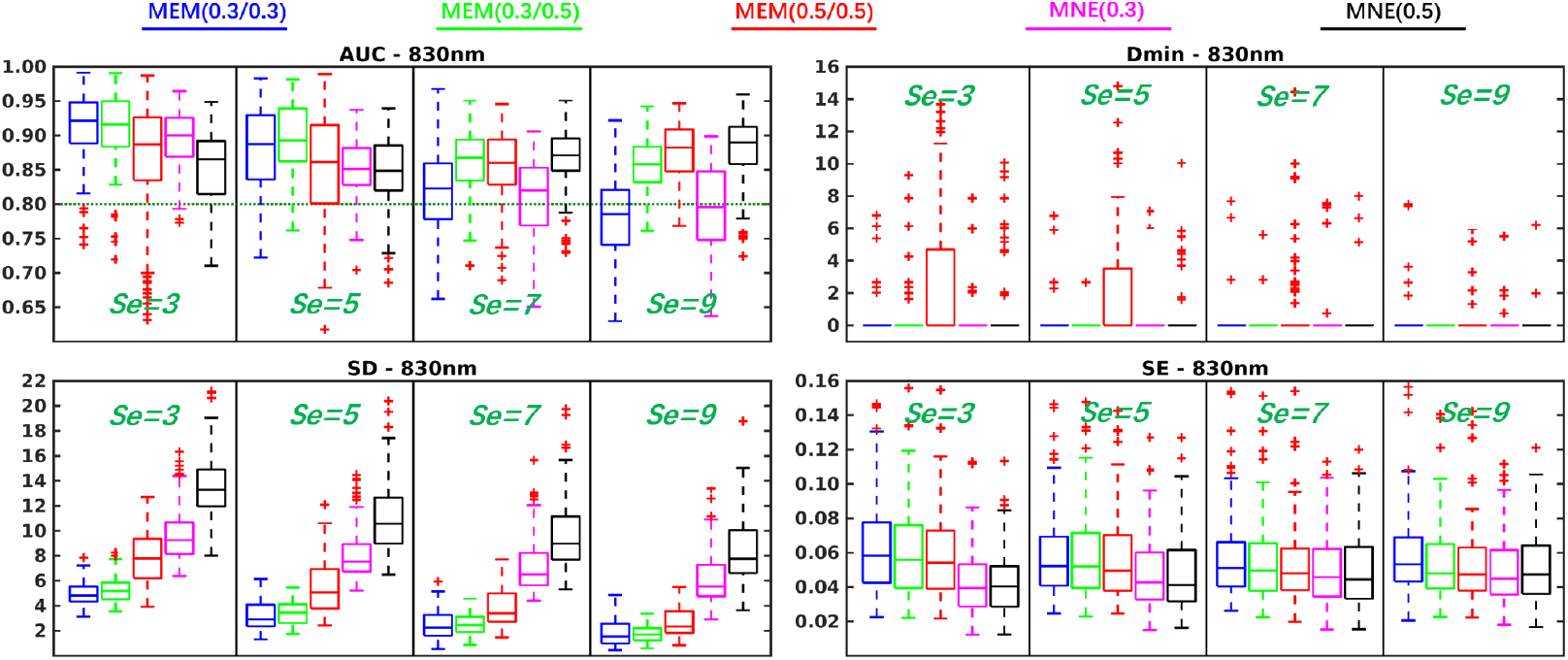
Evaluation of the performances of MEM and MNE using realistic simulations in-volving superficial seeds for different spatial extent (*Se* = 3, 5, 7, 9). Boxplot representation of the distribution of four validation metrics for three depth weighted strategies of MEM and two depth weighted strategies of MNE, namely: MEM(0.3, 0.3) in blue, MEM(0.3, 0.5) in green, MEM(0.5, 0.5) in red, MNE(0.3) in magenta and MNE(0.5) in black. Results were obtained after DOT reconstruction of 830*nm* Δ*OD*.

**Table 1.** Wilcoxon signed rank test results of reconstruction performance comparison of MEM and MNE in superficial seeds case. Median values of paired difference are presented in the table. *p* values were corrected for multiple comparisons using Bonferroni correction, * indicates *p* < 0.01 and ** represents *p* < 0.001. Median of the paired difference of each validation metrics is color coded as follows: green: MEM is significantly better than MNE, red: MNE is significantly better than MEM and gray: non-significance.

| Superficial Seeds |  | Se = 3 |  | Se = 5 |  | Se = 7 |  | Se = 9 |  |
| --- | --- | --- | --- | --- | --- | --- | --- | --- | --- |
|  |  | MNE (0.3) | MNE (0.5) | MNE (0.3) | MNE (0.5) | MNE (0.3) | MNE (0.5) | MNE (0.3) | MNE (0.5) |
| AUC | MEM (0.3, 0.3) | 0.02* | 0.06** | 0.04** | 0.03** | 0.01 | -0.04** | -0.01 | -0.10** |
|  | MEM (0.3, 0.5) | 0.02* | 0.05** | 0.05** | 0.04** | 0.05** | 0.00 | 0.05** | -0.03** |
|  | MEM (0.5, 0.5) | -0.01 | 0.03 | 0.00 | 0.01 | 0.04** | 0.00 | 0.07** | -0.01 |
| Dmin | MEM (0.3, 0.3) | 0.00 | 0.00 | 0.00 | 0.00 | 0.00 | 0.00 | 0.00 | 0.00 |
|  | MEM (0.3, 0.5) | 0.00 | 0.00 | 0.00 | 0.00 | 0.00 | 0.00 | 0.00 | 0.00 |
|  | MEM (0.5, 0.5) | 0.00 | 0.00 | 0.00 | 0.00 | 0.00 | 0.00 | 0.00 | 0.00 |
| SD | MEM (0.3, 0.3) | -4.26** | -8.31** | -4.48** | -7.63** | -4.16** | -6.43** | -3.85** | -6.28** |
|  | MEM (0.3, 0.5) | -3.78** | -8.23** | -4.11** | -7.11** | -3.86** | -6.30** | -3.71** | -6.24** |
|  | MEM (0.5, 0.5) | -1.64** | -5.56** | -2.60** | -4.97** | -2.90** | -4.79** | -2.84** | -5.01** |
| SE | MEM (0.3, 0.3) | 0.02** | 0.02** | 0.01** | 0.01** | 0.01 | 0.01 | 0.01 | 0.01 |
|  | MEM (0.3, 0.5) | 0.02** | 0.02** | 0.01** | 0.01** | 0.01 | 0.01 | 0.00 | 0.00 |
|  | MEM (0.5, 0.5) | 0.02** | 0.02** | 0.01** | 0.01** | 0.00 | 0.00 | 0.00 | 0.00 |

For spatial accuracy, results evaluated using Dmin, we obtained median Dmin values of 0*mm* for all methods, indicating the peak of the reconstructed map, was indeed accurately localized inside the simulated generator. It is worth mentioning that MEM(0.5, 0.5) provided few Dmin values larger than 0*mm* in *Se* = 3 and *Se* = 5 cases, which consisted of superficial and focal generators. Since MEM accurately estimated the spatial extent, more depth weighting considered for MEM(0.5, 0.5) could results in focal and deeper reconstruction, hence resulting in non-zero Dmin values. On the other hand, MNE would over-estimate the size of the underlying generators, therefore resulting in 0*mm* Dmin, but larger SD values in similar conditions.

When considering the general reconstruction accuracy using AUC, for focal generators such as *Se* = 3 and 5, we found significant larger AUC (see Table. 1) for MEM(0.3, 0.3) and MEM(0.3, 0.5) when compared to the most accurate version of MNE, i.e., MNE(0.3). When considering more extended generators, i.e., *Se* = 7 and 9, MEM(0.3, 0.5) and MEM(0.5, 0.5) achieved significantly larger AUC than MNE(0.3). However, the AUC of MNE(0.5) was significantly larger than MEM(0.3, 0.3) for *Se* = 7 as well as significantly larger than MEM(0.3, 0.5) and MEM(0.5, 0.5) for *Se* = 9.

In terms of spatial extent of the estimated generator distribution and the localization error, MEM provided significantly smaller SD values among all the comparisons. Finally, for temporal accuracy of the reconstruction represented by SE, MNE provided significantly lower values, but with a small difference (e.g., 0.01 or 0.02, see results on real data as a reference of this effect size), than MEM among all comparisons when *Se* = 3, 5.

Similar comparison between MEM and MNE were conducted respectively for middle seed simulated generators and deep seed simulated generators. Results were overall reporting similar trends when comparing MEM and MNE methods for middle and deep seeds, and as expected more depth weighting resulted in more accurate reconstructions (described in details in supplementary material, Fig. S2 and Table. S1 for middle seeds, Fig. S3 and Table. S2 for deep seeds).

To further illustrate the performance of MEM and MNE as a function of the depth of the generator, we are presenting some reconstruction results in Fig. 4. Three generators with a spatial extent of *Se* = 5, were selected for this illustration. They were all located around the right “hand knob” area, and were generated from a superficial, middle and deep seed respectively. The first column in Fig. 4 shows the location and the size of the simulated generator, considered as our ground truth. The generator constructed from the superficial seed only covered the corresponding gyrus, whereas the generators constructed from the middle seed, included parts of the sulcus and the gyrus. Finally, when considering the deep seed, the simulated generator covered both walls of the sulcus, extended just a little on both gyri. For superficial case, MEM(0.3, 0.3) and MEM(0.3, 0.5) provided similar performances in term of visual evaluation of the results and quantitative evaluations (*AUC* = 0.96, *Dmin* = 0*mm*, *SD* = 1.94*mm*, 2.15*mm*, *SE* = 0.03). On the other hand, for the same simulations, MNE(0.3) and MNE(0.5) resulted in less accurate reconstructions, spreading too much around the true generator, as confirmed by validation metric, exhibiting notably large SD values (*AUC* = 0.86, 0.89, *Dmin* = 0*mm*, *SD* = 9.84*mm*, 14.63*mm*, *SE* = 0.02). When considering the simulation obtained with the middle seed, MEM(0.3, 0.5) retrieved accurately the gyrus part of the generator but missed the sulcus component, since less depth compensation was considered. When increasing depth sensitivity, MEM(0.5, 0.5) clearly outperformed all other methods, by retrieving both the gyrus and sulcus aspects of the generator, resulting in the largest *AUC* = 0.98 and the lowest *SD* = 2.93*mm*. MNE(0.3) was not able to recover the deepest aspects of the generator, but also exhibited a large spread outside the ground truth area as suggested by a large *SD* = 9.69*mm*. MNE(0.5) was able to retrieve the main generator, but also exhibited a large spatial spread of *SD* = 10.16*mm*. When considering the generators obtained from the deep seed, MNE(0.3) only reconstructed part of gyrus, missing completely the main sulcus aspect of the generator, resulting in low AUC of 0.57 and large SD of 10.34*mm*. MEM(0.3, 0.5) was not able to recover the deepest aspects of the sulcus, but reconstructed accurately the sulci walls, resulting in an AUC of 0.89 and a SD of 2.71*mm*. MEM(0.5, 0.5) recovered the deep simulated generator very accurately, as demonstrated by the excellent scores (*AUC* = 0.97, *SD* = 2.11*mm*) when compared to MNE(0.5). For those three simulations, all methods recovered the underlying time course of the activity with similar accuracy (i.e., similar SE values). In supplementary material, we added Video.1, illustrating the behavior of all the simulations and all methods, following the same layout provided in Fig. 4.

**Fig. 4.**
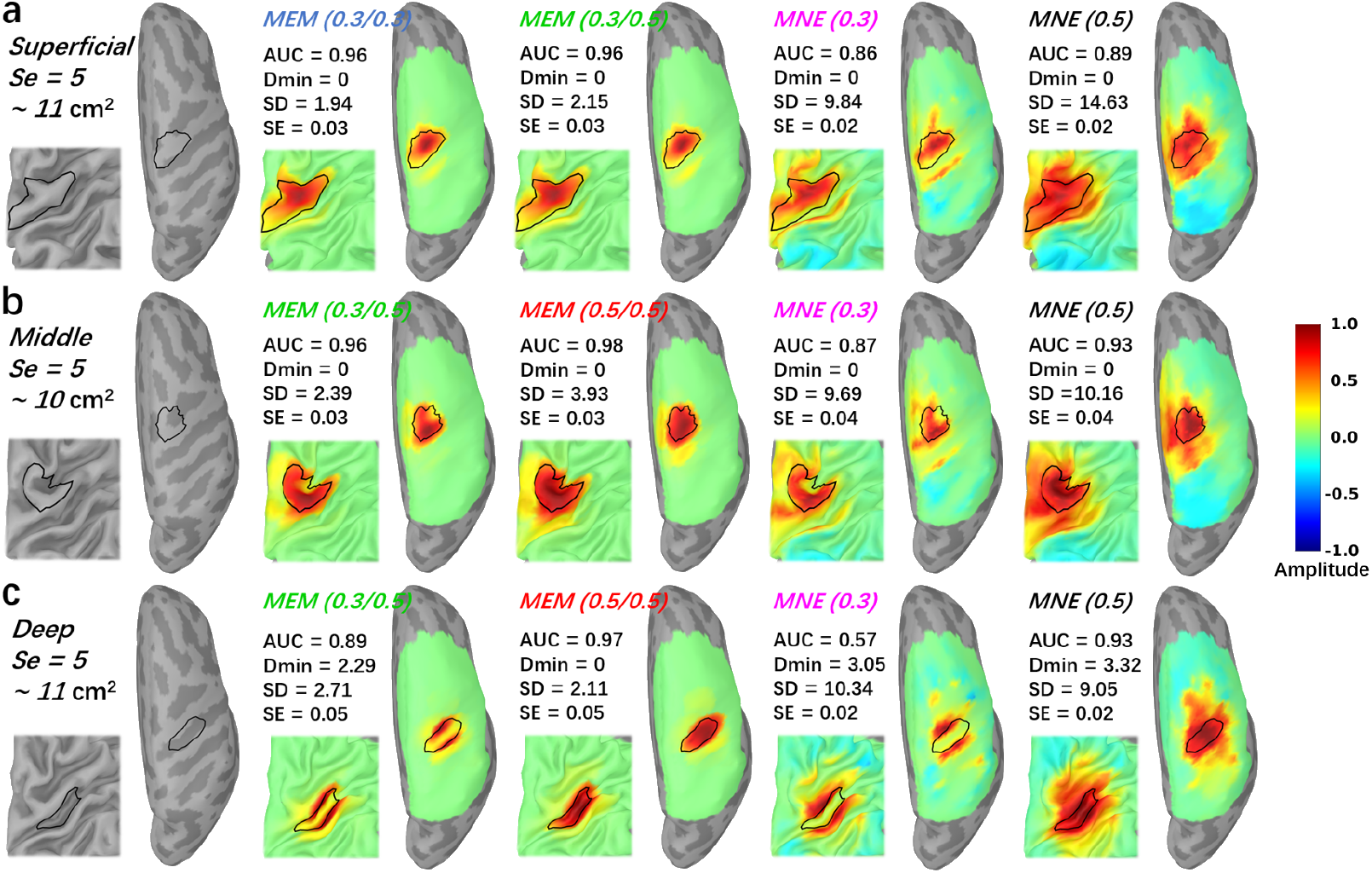
Comparisons of the reconstruction maps using MEM and MNE in realistic simulations. Three theoretical regions with spatial extent *Se* = 5 (11*cm*^2^) were selected near the hand knob at different depth. The first column presents the locations and the size of the generator along the cortical surface. (a) Superficial seed case with reconstructed maps reconstructed using all MEM and MNE implementations considered in this study. (b) Middle seed case with reconstructed maps reconstructed using all MEM and MNE implementations considered in this study. (c) Deep seed case with reconstructed maps reconstructed using all MEM and MNE implementations considered in this study. 20% inflated and zoomed maps are presented on the left corner of each figure. 100% inflated right hemisphere are presented on the right side. All the maps were normalized by their own global maximum and no threshold was applied.

Note that for this quantitative evaluation of fNIRS reconstruction methods using realistic simulation framework, we considered fNIRS data at only one wavelength (830*nm*). Using single wavelength in the context simulation based evaluation is a common procedure in DOT literature (Zhan et al., 2012; Dehghani et al., 2009; White and Culver, 2010; Okawa et al., 2011; Tremblay et al., 2018; Shimokawa et al., 2012, 2013), since we may expect overall similar performances for 685*nm* wavelength under the same SNR level.

### 3.2. Effects of depth weighting on the reconstructed generator as a function of the depth and size of the simulated generators

To summarize the effects of depth weighting in 3D fNIRS reconstructions, we further investigated the validation metrics, AUC, SD and SE, as a function of depth and size of the simulated generators. Dmin was not included due to the fact that we did not find clear differences among methods throughout all simulation parameters from previous results. In the top row of Fig. 5, 250 generators created from all 250 seeds with a spatial extent of *Se* = 5 were selected to demonstrate the performance of different versions of depth weighting as a function of the average depth of the generator. Whereas in the bottom row of Fig. 5, we considered 400 generators constructed from all 100 superficial seeds with 4 different spatial extents of *Se* = 3, 5, 7, 9, to illustrate the performance of different versions of depth weighting as a function of the size of the generator. According to AUC, depth weighting was indeed necessary for all methods when the generator moved to deeper regions (> 2*cm*) as well as when the size was larger than 20*cm*^2^. Moreover, any version of MEM always exhibited clearly less false positives, as indicated by lower SD values, than all of MNE versions, whatever was the depth or the size of the underlying generator. We found no clear trend and difference of temporal accuracy among methods when reconstructing generators of different depths and sizes.

**Fig. 5.**
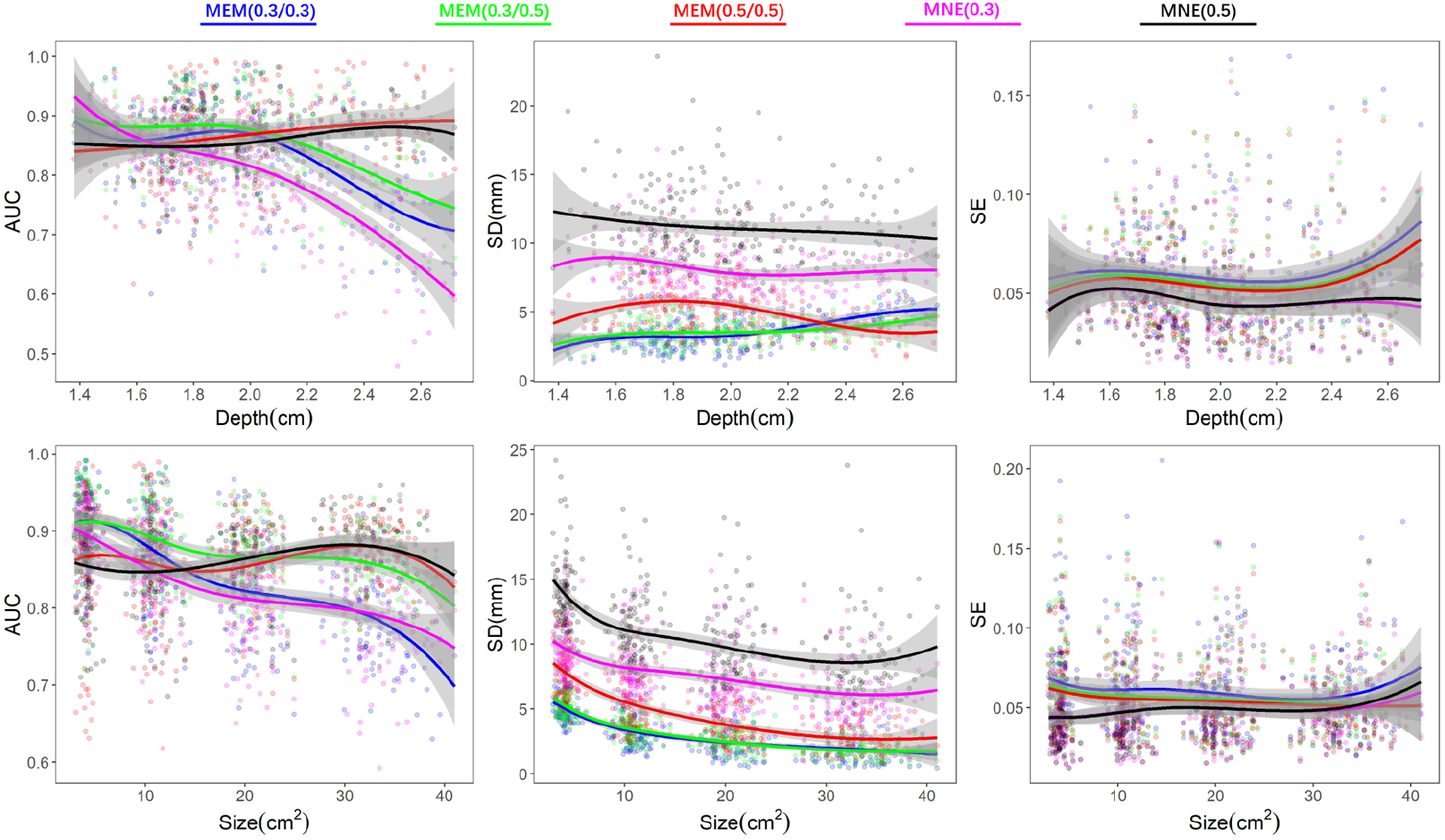
Effects of depth weighting on the depth and size of the simulated generators. First row demonstrates the validation matrices, AUC, SD and SE, as a function of depth of generators. We selected 250 generators created from all 250 seeds with a spatial extent of *SD* = 5. Depth was calculated by the average of minimum Euclidean distance from each vertex, within each generator, to the head surface. Second row demonstrates the validation matrices, AUC, SD and SE, as a function of size of generators. Involving 400 generators which constructed from 100 superficial seeds with 4 different spatial extend of *Se* = 3, 5, 7, 9. Line fittings were performed via a 4 knots spline function to estimate the smoothed trend and the shade areas represent 95% confident interval. Color coded points represent the values of validation matrices of all involved generators.

### 3.3. Robustness of 3D reconstructions to the noise level

All previous investigations were obtained from simulations obtained with a SNR of 5, in this section we compared the effect of the SNR level in Fig. 6, on depth weighted versions of MNE and MEM, for superficial seeds only and generators of spatial extent *Se* = 5. We only compared MEM(0.3, 0.5) and MNE(0.5) considering the observation from previous results that these two methods were overall exhibiting best performances in this condition. Regarding Dmin, paired differences were not significant but MNE exhibited more Dmin values above 0*mm* than MEM at all SNR levels, suggesting that MNE often missed the main generators while MEM was more accurate in reconstructing the maximum of activity within the simulated generator. Regarding AUC, MEM(0.3, 0.5) exhibited values higher than 0.8 at all SNR levels, whereas MNE(0.5) failed to recover accurately the generator for *SNR* = 1. Besides, in Table. 2, we found that difference of AUC between MEM and MNE increased when SNR level decreased, suggesting the good robustness of MEM when decreasing the SNR level. The difference of SD also increased when SNR levels decreased. Indeed, MEM exhibited stable SD values among most SNR levels (except *SNR* = 1), whereas for MNE SD values were highly influenced by the SNR level. Finally, for both methods, decreasing SNR levels resulted in less accurate time course estimation (SE increased), slightly more for MEM when compared to MNE.

**Fig. 6.**
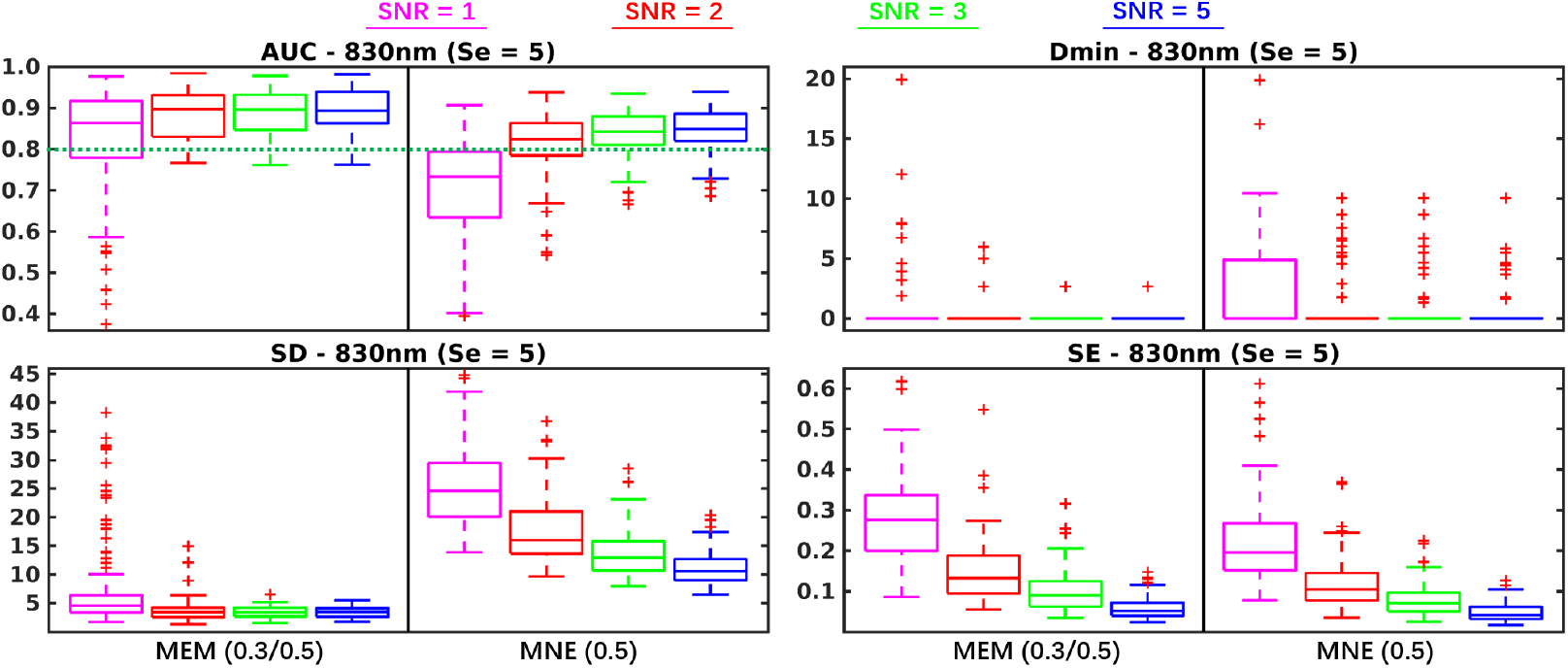
Evaluation of the performances of MEM and MNE at four different SNR levels. Boxplot representation of the distribution of four validation metrics for MEM(0.3, 0.5) and MNE(0.5) involving superficial seeds with spatial extent *Se* = 5. SNR levels (*SNR* = 1, 2,3, 5) are represented using different colors.

**Table. 2.** Reconstruction performance comparison of MEM and MNE with different SNR levels. Median of paired difference of validation metric (i.e. AUC, Dmin, SD and SE) values of *Se* = 5 are presented in the table following the SNR increase from 1 to 5. ** indicates corrected *p* < 0.001.

| $Se = 5$ ($\sim 11 \text{ cm}^2$) | | SNR = 1 | SNR = 2 | SNR = 3 | SNR = 5 |
| --- | --- | --- | --- | --- | --- |
|  |  | MNE (0.5) | MNE (0.5) | MNE (0.5) | MNE (0.5) |
| AUC | MEM (0.3, 0.5) | 0.14** | 0.07** | 0.05** | 0.04** |
| Dmin | MEM (0.3, 0.5) | 0.00 | 0.00 | 0.00 | 0.00 |
| SD | MEM (0.3, 0.5) | -17.63** | -12.40** | -9.22** | -7.11** |
| SE | MEM (0.3, 0.5) | 0.05** | 0.03** | 0.02** | 0.01** |

### 3.4. Evaluation of MEM and MNE on real fNIRS data

For all finger tapping fNIRS data considered in our evaluations, two wavelength (i.e., 685*nm* and 830*nm*) were reconstructed first and then converted to HbO/HbR concentration changes along cortical surface using specific absorption coefficients. All the processes from fNIRS preprocessing to 3D reconstruction were completed in Brainstorm (Tadel et al., 2011) using the NIRSTORM plugin developed by our team (https://github.com/Nirstorm). For full double density montage (montage 1), reconstructed HbR amplitudes were reversed to positive phase and normalized to their own global maximum, to facilitate comparisons. In Fig. 7.a, we showed the re-constructed HbR maps at the peak of the time course (i.e., 31s) for MEM and MNE by considering the 4 depth weighted versions, previously evaluated, i.e., MEM(0.3, 0.3), MEM(0.3, 0.5), MNE(0.3) and MNE(0.5). The two depth weighted versions of MEM clearly localized well the “hand knob” region, while exhibiting very little false positives in its surrounding. On the other hand, both depth weighted version of MNE clearly overestimated the size of the hand knob region and were also exhibiting some distant possibly spurious activity. The fMRI Z-map obtained during the corresponding fMRI task is presented on Fig. 7.b, after projection of the volume Z-map on the cortical surface. Fig. 7.c showed the time courses within the region of interest representing the “hand knob”. Each curve represents the reconstructed time course of one vertex of the hand knob region and the amplitude were normalized by the peak value within the whole region.

**Fig. 7.**
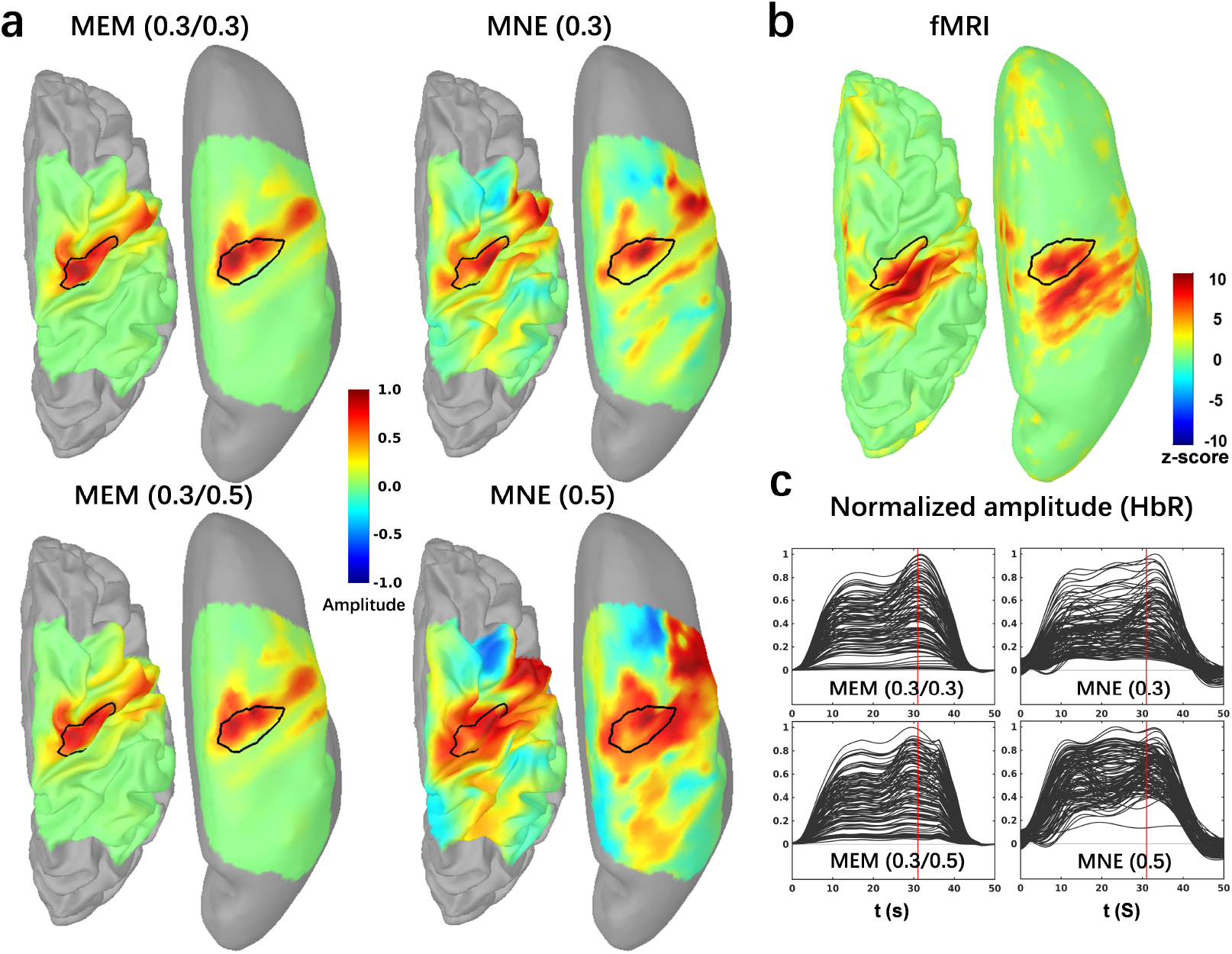
Application of MEM versus MNE reconstruction of HbR during a finger tapping task on one healthy subject. (a) Reconstructed maps of HbR (e.g. 20% inflation on the left and 100% inflation on the right side.) from MEM and MNE with different depth compensations. Each map was normalized by its own global maximum. (b) fMRI Z-map results projected along the cortical surface. (c) Reconstructed time courses of HbR within the hand knob region from MEM and MNE. Note that the hand knob region, represented by the black profile, was also matched well with the mean cluster of fMRI activation map on primary motor cortex. No statistical threshold was applied on fNIRS reconstructions.

Results obtained on 5 subjects for acquisition involving personalized optimal fNIRS montage (montage 2) and corresponding fNIRS reconstructions are presented in Fig. 8. For every subject, fMRI Z-maps are presented along the left hemisphere only and thresholded at *Z* > 3.1 (*p* < 0.01, corrected us-ing Gaussian random field theory), The most significant fMRI cluster along M1 and S1 was delineated using a black profile. Reconstruction maps at the corresponding HbO/HbR peaks are then presented. Similar accuracy between MEM and MNE, with good overlap with fMRI results, were found for subjects 4 and 5, while MNE was overestimating the spatial extent of the generator. For subject 1, 2 and 3, MNE exhibited poor spatial correspondence with fMRI results. Averaged reconstructed time courses within the fMRI main cluster region are shown with standard deviation as the error bar. Comparing to simulations results, MEM exhibited overall very similar time course estimations than MNE in all cases. Considering the task duration was 10s, the reconstructed peak timing of HbO/HbR appeared accurately within the range of 10*s* to 20*s*.

**Fig. 8.**
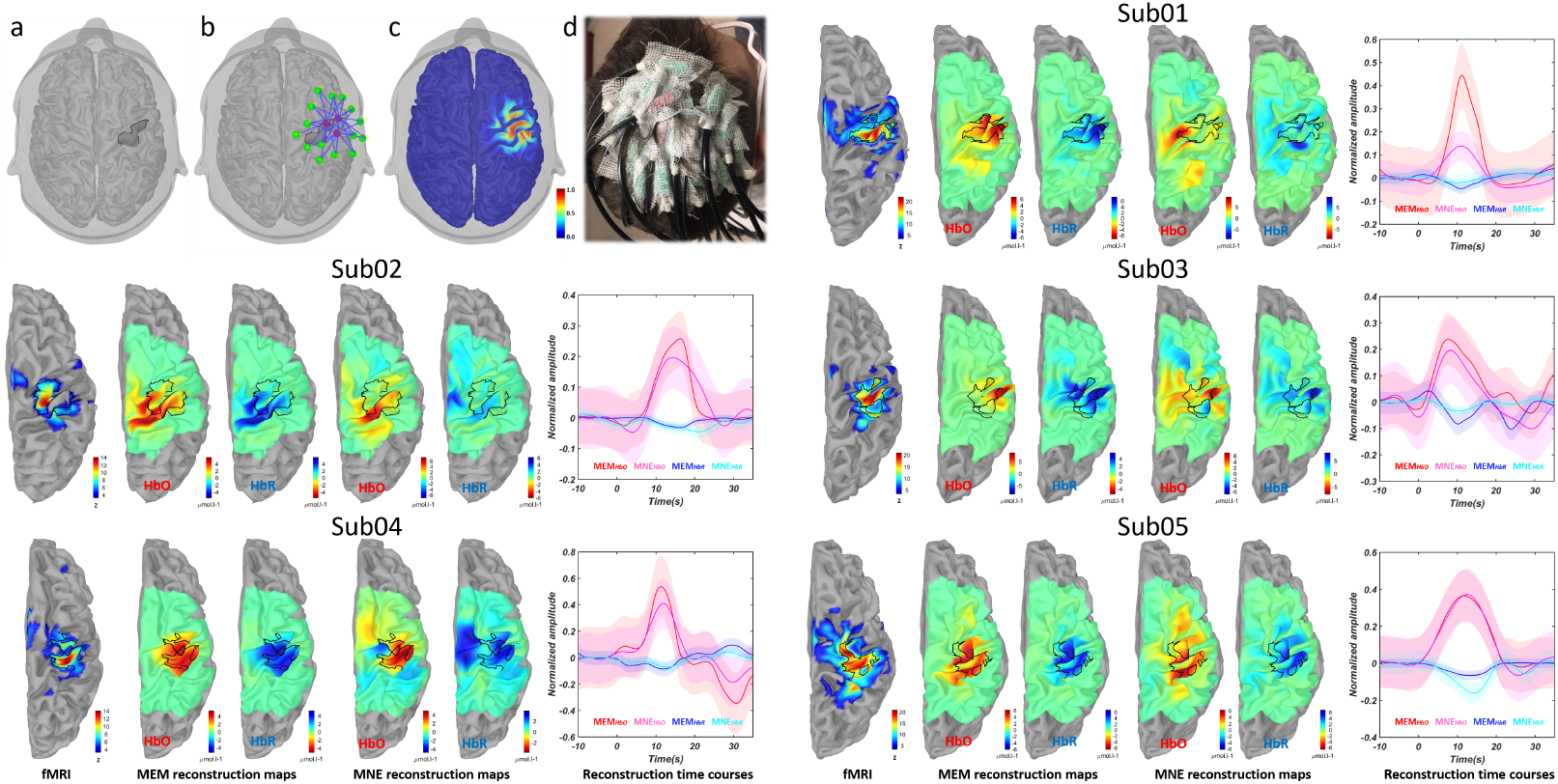
Personalized fNIRS montage and comparisons between MEM and MNE recon-structions with respect to fMRI Z-map at individual level. a) the region of interest defined as the hand knob, b) optimal montage targeting the ROI consisting 3 sources (red) and 15 detectors(green) and one proximity (in the center of sources not shown), c) normalized sensitivity profile of the optimal montage which calculated as the sum of all channels sensitivity along the cortical surface, d) optimal montage glued on the scalp of the one subject, using collodion. fMRI Z-map of each subject during finger tapping task (threshold with *Z* > 3.1, Bonferroni corrected), black profile represents the main cluster along M1 and S1. MEM reconstruction maps at the corresponding HbO/HbR peak times, using depth weighted option 0.3, 0.3. MNE reconstruction maps, at the corresponding HbO/HbR peak times, using depth weighted option 0.3. Reconstructed time courses within the black profile, solid lines represent the main time courses and the shade areas represent standard deviation within the region of interest. Reconstructed time courses were normalized by the maximum amplitude, for each method respectively, before averaging.

## 4. Discussion

### 4.1. Spatial accuracy of 3D fNIRS reconstruction using MEM

In the present study, we first adapted the MEM framework in the context of 3D fNIRS reconstruction and extensively validated its performance. The spatial performance of reconstructions can be considered in two aspects, 1) correctly localizing the peak of the reconstructed map close enough to the ground truth area, 2) accurately recovering the spatial extent of the generator. According to our comprehensive evaluations of the proposed depth-weighted implementations of MEM and MNE methods, accurate localization was overall not difficult to achieve as suggested by our results using Dmin metric. Almost all methods provided median value of Dmin to be 0*mm* in all simulation conditions except for the lowest *SNR* = 1 condition where more localization error was found. On the other hand, recovering the actual spatial extent of the underlying generator is actually the most challenging task in fNIRS reconstruction. When considering the results of MNE on both realistic simulations and real finger tapping tasks, either from visual inspection (Fig. 4, Fig. 7 and Fig. 8) or quantitative evaluation by SD (Fig. 3, Table. 1 and supplementary section S2), we found that MNE overall reconstructed well the main generator but largely overestimated the size of the underlying generator. MEM was specifically developed, in the context of EEG/MEG source imaging, as a method able to recover the spatial extent of the underlying generators, which has been proved not to be the case for MNE-based approaches (Chowdhury et al., 2013, 2016; Grova et al., 2016; Hedrich et al., 2017; Pellegrino et al., 2020). A recent review (Sohrabpour and He, 2021) in the context of EEG/MEG source imaging has also demonstrated that the Bayesian approach with sparsity constraints is required to accurately estimate the spatial extent. These important properties of MEM was successfully demonstrated in our results on fNIRS reconstructions. These excellent performances were reliable for different sizes and depths of simulated generators, and for real finger tapping fNIRS data as well.

### 4.2. Importance of depth weighting in 3D fNIRS reconstruction

Biophysics models of light diffusion in living tissue are clearly demonstrating that, at all source-detector separations, light sensitivity decreases exponentially with depth (Strangman et al., 2013). The general solution to grant the ability of depth sensitivity compensation in fNIRS reconstruction is to introduce depth weighting during the reconstruction. In this study, we investigated the impact of depth weighting effects on fNIRS reconstruction, as a function of the location and the spatial extent of the underlying generators. Our results are showing that when considering little or no depth weighting (*ω* = 0.0 and 0.1) only most superficial generators along the gyral crown were accurately reconstructed missing the deepest parts, therefore resulting in low AUC values. On the other hand, larger depth weighted values, *ω* = 0.7 and 0.9, would bias too much the importance of deep generators and consequently, the most superficial aspects of the underlying generators were not recovered. According to our detailed evaluation on MNE reported in Fig. S1, depth weighted values of *ω* = 0.3 and 0.5 were considered as good candidates to offer an ideal trade off. As expected, MNE(0.5) reported larger spatial dispersion around the true generator, than MNE(0.3). Depth weighting was also important when recovering more extended generators (> 20*cm*^2^, Fig. 5), for both MNE and MEM, since those extended generators were actually involving both superficial and deep regions.

### 4.3. Implementation of depth weighting strategy within the MEM framework

In this study, we are proposing for the first time a depth weighting strategy within the MEM framework, by introducing two parameters: *ω*_1_ acting on scaling the source covariance matrix, and *ω*_2_ tuning the initialization of the reference for MEM. When compared to depth weighted MNE, the MEM framework demonstrated its ability to reconstruct, different depth of focal generators as well as larger size generators, exhibiting excellent accuracy and few false positives (see Fig. 5). When considering deeper focal generators (*depth* > 2*cm*), MEM(0.5,0.5) clearly outperformed all other methods (see AUC and SD values in Fig 5). In summary, for a large range of depths and spatial extents of the underlying generators, MEM methods exhibited accurate results (large AUC values) and less false positives (lower SD values) when compared to MNE methods.

In practice, we would suggest to consider either *ω*_2_ = 0.3 or 0.5 for the initialization of MEM in all cases and only tune *ω*_1_. This is due to the fact that MNE(0.3 or 0.5) provided a generally good reconstruction with larger true positive rate in most scenarios, therefore providing MEM an accurate reference model (*dν*(*x*)) to start with. Even when considering the most focal simulated generators (*Se* = 3) case (see Fig. 3, Table. 1 and Fig. 5), MEM(0.3, 0.3) and MEM(0.3, 0.5) were actually exhibiting very similar performances. Our proposed suggestion to tune *ω*_1_ and *ω*_2_ parameters was actually further confirmed when considered results obtained from real data. For both montages, MEM(0.3, 0.3) results in excellent spatial agreement with fMRI Z-maps.

Note that depth weighting was also considered in DOT studies using MNE (Culver et al., 2003; Zeff et al., 2007; Dehghani et al., 2009; White et al., 2009; Eggebrecht et al., 2012, 2014) and a hierarchical Bayesian DOT algorithm (Shimokawa et al., 2012, 2013; Yamashita et al., 2016). A spatially-variant regularization parameter *β* was added to a diagonal regularization matrix featuring the sensitivity of every generator (forward model), and the value of *β* was tuned according to the sensitivity value of a certain depth. In practice, this strategy would result in similar depth compensation as ours, but we preferred the depth weighting parameter *ω* which mapped the amount of compensation from 0 to 1 (as described in Eq. 3) for easier interpretation and comparison. This is also a standard procedure introduced in EEG/MEG source localization studies (Fuchs et al., 1999; Lin et al., 2006). Finally, using the depth weighted MNE solution as the prior is a common consideration in Hierarchical Bayesian framework based fNIRS reconstructions (Shimokawa et al., 2012, 2013; Yamashita et al., 2016)

### 4.4. Temporal accuracy of 3D fNIRS reconstruction using MEM

Another important contribution of this study was that we improved the temporal accuracy time courses estimated within the MEM framework, resulting in similar temporal accuracy the one obtained with MNE. For instance, the largest significant SE difference between MEM and MNE was only 0.02 for *Se* = 3 and 0.01 for *Se* = 5. Corresponding time course estimations are also reported for MEM and MNE in real data (Fig. 7 and Fig. 8), suggesting again very similar performances. For instance, SE between MEM and MNE HbO time course was estimated as 0.02 for *Sub*05 in Fig. 8. Moreover, we found no significant SE differences between MEM and MNE for more extended generators (Se = 7,9). These findings are important considering that MNE is just a linear projection therefore the shape of the reconstruction will directly depend on the averaged signal at the channel level. On the other hand, MEM is a nonlinear technique, applied at every time sample, which is not optimized for the estimation of resulting time courses.

### 4.5. Robustness of fNIRS reconstructions to the noise level

To further investigate the effects of the amount of realistic noise in our reconstructions on both reconstruction methods, we performed the comparisons along 4 different SNR levels, i.e., *SNR* = 1, 2, 3, 5. As shown in Fig. 6 and Table. 2, we found that MEM was overall more robust than MNE when dealing with simulated signals at lower SNR levels. This is actually a very important result since when reconstructing HbO/HbR responses, one has to consider at least two ΔOD of two different wavelengths exhibiting different SNR levels. For the simulation results, we reported reconstruction results obtained from 830*nm* data, whereas when considering real data (Fig. 7 and Fig. 8), we had to convert the reconstruction absorption changes at 685*nm* and 830*nm* into HbO/HbR concentration changes. Therefore, our final results were influenced by the SNR of all involved wavelengths.

fNIRS is inherently sensitive to inter-subject variability (Novi et al., 2020), as also suggested in our application on real data presented in Fig. 8. Data from *Sub*05 were exhibiting a good SNR level and therefore both MEM and MNE reconstructed accurately the main cluster of the activation, while MNE presented more spatial spread and false positive activation outside the fMRI ROI. When considering subjects for whom we obtained lower SNR data, e.g., *Sub*02 and *Sub*03, MEM still recovered an activation map similar to fMRI map. In those cases, MNE not only reported suspicious activation pattern but also incorrectly reconstruct the peak amplitude outside the fMRI ROI. Our results suggesting MEM robustness in low SNR conditions for DOT are actually aligned with similar findings suggested for EEG/MEG source imaging, when considering source localization of single trial data (Chowdhury et al., 2018; Aydin et al., 2020).

### 4.6. Comprehensive evaluation and comparison of the reconstruction performance using MEM and MNE

To perform a detailed evaluation of our proposed fNIRS reconstructions methods, we developed a fully controlled simulation environment, similar to the one proposed by our team to validate EEG/MEG source localization methods (Chowdhury et al., 2013, 2016; Hedrich et al., 2017). The fNIRS resting state data, acquired by the same montage (montage1) and underwent the same preprocessing as conducted for the real data, was added to the simulated true hemodyanmic response for each channel. Indeed such environment provided us access to a ground truth, which is not possible when considering real fNIRS data set. Previous studies validated tomography results (Eggebrecht et al., 2014; Yamashita et al., 2016) by comparing with fMRI activation map which can indeed be considered as a ground truth, but only for well controlled and reliable paradigms. Since fMRI also measures a signal of hemodynamic origin, it is reasonable to check the concordance between fMRI results and DOT reconstructions. Therefore, as preliminary illustrations, we also compared our MEM and MNE results to fMRI Z-maps obtained during finger tapping tasks on 6 healthy participants, suggesting overall excellent performances of MEM when compared to MNE. Further quantitative comparison between fMRI and fNIRS 3D reconstruction, was out of the scope of this paper and will be considered in future studies.

### 4.7. Sampling size of fNIRS reconstructions

As opposed to several other fNIRS tomography studies that reconstruct fNIRS responses within a 3D volume space, here we proposed to use the mid-cortical surface as anatomical constraint to guide DOT reconstruction. However, the maximum spatial resolution of our surface based reconstruction was similar to the volume based one. Indeed, DOT reconstruction within a volume space usually down-sampled light sensitivity maps to either 2 × 2 × 2 mm^3^ (Eggebrecht et al., 2014), 3 × 3 × 3 mm^3^ (Eggebrecht et al., 2012) or 4 × 4 × 4 mm^3^ (Yamashita et al., 2016) matrices, resulting in the downsampled voxel volume ranging from 8mm^3^ to 64mm^3^. In our case, when projecting from volume space into cortical surface space, a unique set of voxels were assigned to each vertex along the cortical surface according to the Voronoi based projection method (Grova et al., 2006). Considering the mid-surface resolution (i.e., 25, 000 vertices) used in this study, the average volume of a Voronoi cell was 25mm^3^, which falls in the above volume range. Therefore, we can conclude that both volume-based and surface-based fNIRS reconstructions as implemented here would result in similar sampling of the reconstruction space.

### 4.8. fNIRS montage for 3D reconstructions

In previous reported studies (Zeff et al., 2007; White and Culver, 2010; Zhan et al., 2012; Eggebrecht et al., 2012, 2014), a high density montage was considered which was proved to be able to provide high spatial resolution and robustness to low SNR conditions (White and Culver, 2010). In the present study, we first considered a full double density montage, as proposed in (Kawaguchi et al., 2007), to generate realistic simulations, and then analyzed finger tapping results on real data acquired on one subject. Double density montages have been involved in several inverse modelling studies such as (Kawaguchi et al., 2004; Sakakibara et al., 2016; Machado et al., 2018). We also illustrated, in 5 other subjects, MEM performance when considering real data set acquired by optimal montages, exhibiting a large amount of local spatial overlap between channels. In this case, probe design was optimized to maximize the sensitivity to the hand knob ROI, while also ensuring sufficient spatial overlap between sensors (e.g., at least 13 detectors had to construct channels with each of the three sources, and the channel distance was ranging from 2*cm* to 4.5*cm*, see Fig. 8a). We have previously demonstrated in Machado et al., 2018 that even if high density montages are usually considered as a gold standard for DOT reconstruction, personalized optimal montages (Machado et al., 2014, 2018, 2021) have ability to allow accurate reconstructions along the cortical surface. Finally, evaluating the performance of MEM when considering high density fNIRS montage would be of great interest but was out of the scope of this present study.

### 4.9. Availability of the proposed MEM framework

Several software packages have been proposed to provide fNIRS reconstruction pipelines, as for instance NeuroDOT (Eggebrecht et al., 2014, 2019), AtlasViewer(Aasted et al., 2015) and fNIRS-SPM(Ye et al., 2009). To ensure an easy access of our MEM methodology to the fNIRS community, we developed and released a fNIRS processing toolbox - NIRSTORM (https://github.com/Nirstorm), as a plugin of Brainstorm software (Tadel et al., 2011), which is a renown software package dedicated for EEG/MEG analysis and source imaging. Our package NIRSTORM offers standard preprocessing, analysis and visualization as well as more advanced features such as personalized optimal montage design, access to forward model estimation using MCXlab(Fang and Boas, 2009; Yu et al., 2018) and the MNE and MEM implementations considered in this study.

### 4.10. Limitations and Perspectives

Previously, Tremblay et al., 2018 had comprehensively compared a variety of fNIRS reconstruction methods using large number of realistic simulations. Since introducing MEM was our main goal of this study, we did not consider such wide range of methodological comparisons. We decided to carefully compare MEM with MNE since MNE remains the main method considered for DOT, and is available in several software packages. As suggested in Tremblay et al., 2018, DOT reconstruction methods based on Tikhonov regularization, such as least square regularization in MNE, usually allow great sensitivity, but performed poorly in term of spatial extent - largely overestimating the size of the underlying generator. On the other hand, L1-based regularization (Süzen et al., 2010; Okawa et al., 2011; Kavuri et al., 2012; Prakash et al., 2014) could achieve more focal solutions with high specificity but much lower sensitivity. As demonstrated in our results, the proposed MEM framework allows reaching good sensitivity and accurate reconstruction of the spatial extent of the underlying generator. Bayesian model averaging (BMA) originally proposed for EEG source imaging by Trujillo-Barreto et al., 2004, also allows accurate DOT reconstructions with less false positives when compared to MNE. Similarly, we carefully compared MEM to Bayesian multiple priors approaches in Chowdhury et al., 2013 in the context of MEG source imaging. Comparing MEM with more advanced DOT reconstruction methods, including also the one proposed by Yamashita et al., 2016, would be of great interest but was out of the scope of this study.

Overall the main advantage of the MEM framework is its flexibility. Since the core structure of the MEM framework is to provide a unique reconstruction map by maximizing the entropy relative to a reference source distribution, one could implement its own reference for specific usage. For instance, as considered in the present study, the reference distribution considered the depth weighting MNE solution and spatial smoothing to inform our prior model for MEM. Note that in this study we applied MEM independently for the two wavelengths and then calculated HbO/HbR concentration changes after reconstruction, whereas one could directly solve HbO/HbR concentration changes along with reconstructions. Such procedure has been suggested by Li et al., 2004, by incorporating signals from the two wavelength within the same DOT reconstruction model. In the future, the MEM framework would allow to easily implement such a fusion model, as suggested by Chowdhury et al., 2015 in the context of MEG/EEG fusion algorithms. We have shown that MEM-based EEG/MEG fusion allows higher reliability in the source imaging results (Chowdhury et al., 2018), we will consider such an approach to estimate directly HbO/HbR fluctuations from the two wavelengths signals.

Finally, considering the main contribution of this study was to introduce the MEM framework for 3D fNIRS reconstruction, we decided to first carefully evaluate the performance of MEM, using well controlled realistic simulations. We also included few real data set reconstructions to illustrate the performance of the MEM reconstruction, whereas quantitative evaluation of MEM reconstructions on larger database will be considered in our future investigations.

## 5. Conclusion

In this study, we introduced a new fNIRS reconstruction method entitled Maximum Entropy on the Mean (MEM). We first implemented depth weighting into MEM framework and improved its temporal accuracy. To carefully validate the method, we applied a large number (*n* = 4000) of realistic simulations with various spatial extents and depths. We also evaluated the robustness of the method when dealing with low SNR signals. The comparison of the proposed method with the widely used depth weighted MNE was performed by applying four different quantification validation metrics. We found that MEM framework provided accurate and robust reconstruction results, relatively stable for a large range of spatial extents, depths and SNRs of the underlying generator. Moreover, we implemented the proposed method into a new fNIRS processing plugin - NIRSTORM in Brainstorm software to provide the access of the method to users for applications, validations and comparisons.

## Acknowledgments

This work was supported by the Natural Sciences and Engineering Research Council of Canada Discovery Grant Program (CG and JML) and an operating grant from the Canadian Institutes for Health Research (CIHR MOP 133619 (CG)). fNIRS equipment was acquired using grants from NSERC Research Tools and Instrumentation Program and the Canadian Foundation for Innovation (CG). ZC is funded by the Fonds de recherche du Qubec Sante (FRQS) Doctoral Training Scholarship and the PERFORM Graduate Scholarship in Preventive Health Research. GP is funded by Strauss Canada Foundation.

## Data availability

The original raw data supporting the findings of this study are available upon reasonable request to the corresponding authors.

## Appendix A. Supplementary material

Supplementary material associated with this article can be found at the end of this manuscript.

## Supplementary material

### S1. Validation metrics

Here is a detailed description of the four validation metrics considered in our evaluation. Except the shape error (SE), other metrics were all calculated at the time instant *τ* when the simulated Δ*OD* time course reached its peak value (e.g. 12.2s after onset).

**Area Under the Receiver Operating Characteristic (ROC) curve (AUC)** was used to assess overall detection accuracy of the reconstruction methods. We used a specific version of AUC that has been proposed in (Grova et al., 2006) in order not to bias results towards false positives. In further details, ROC curves were generated by plotting the sensibility of the detection as a function of 1-specificity, while thresholding the normalized reconstruction map from 0 to 1 with a certain step value. In the context of source reconstruction, especially when the generator is focal, the region of true positive is usually much smaller than the region of true negative, whereas non-biased AUC evaluation would require to sample the same amount of active and inactive generators. To overcome this possible bias, we considered a ROC evaluation using the same number of active and inactive generators that were randomly sampled within two different regions: 1) *AUC_close_*: inactive generators were sampled within the immediate spatial neighborhood of the ground truth; and 2) *AUC_far_*: inactive generators were sampled within the local maxima of the reconstructed activity located far from the ground truth. The final AUC was then the average of *AUC_close_* and *AUC_far_*.

**Minimum geodesic distance (Dmin)** was represented by the geodesic distance, following the circumvolutions of the cortical surface, of the vertex that exhibited maximum of reconstructed activity to the border of the ‘generator’. It should be 0 when the peak of the reconstruction map was located inside the simulated cortical region.

**Spatial Dispersion (SD)** assessed the spatial spread of the estimated ‘generator’ distribution and the localization error using Eq. S1. The ideal value (i.e. *SD* = 0*mm*), was achieved when no activation was reconstructed outside the theoretical ‘generator’. The larger the SD was, the more spatially spread were the reconstructed maps.

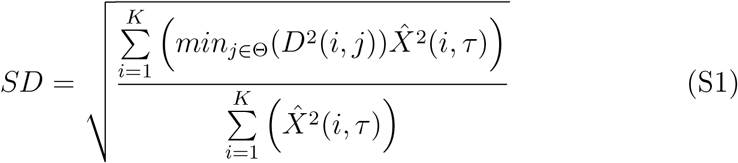

where *min_j∈Θ_*(*D*^2^(*i, j*)) is the minimum Euclidean distance between the vertex *i* to the vertex *j* which is located inside the simulated ‘generator’ (Θ). 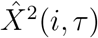 is the power of the amplitude of reconstructed time course on vertex *i* at time *τ*. *K* is the total number of vertices within the reconstruction field of view.

**Shape error(SE)** evaluated the temporal accuracy of the reconstruction. Reconstructed time courses within the simulation ‘generator’ were averaged and normalized. The root mean square of the difference between this time course and the normalized theoretical time course was estimated and denoted as SE in Eq. S2 as introduced in (Chowdhury et al., 2013)

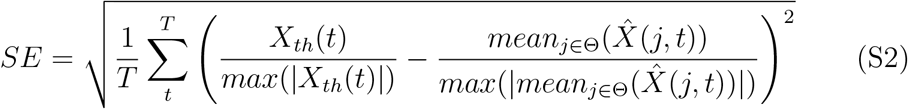

where *T* is length of the time course. *X_th_*(*t*) is the theoretical time course of the simulation. 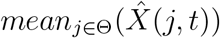 is the averaged mean of the reconstructed time courses within the ‘generator’.

### S2. Effects of depth weighting on MNE

We first investigated the effects of depth weighting factor *ω*_2_ selection for depth weighted MNE. To do so, we evaluated spatial and temporal performances of DOT reconstruction. As presented in Fig. S1, we compared depth weighted MNE using depth weighting factors *ω*_2_ = 0, 0.1, 0.3, 0.5, 0.7,0.9 in superficial seeds case. In general, *ω*_2_ = 0.3 and 0.5 provided overall the most accurate results (i.e. median *AUC* > 0.8 and *Dmin* = 0*mm*). For focal generators(i.e. *Se* = 3, 5), *ω*_2_ = 0.3 performed better than *ω*_2_ = 0.5 considering it was providing significantly lower SD. However, in extended generators (i.e. *Se* = 7,9), reconstructions with *ω*_2_ = 0.5 were exhibiting more accurate results, consisting in significantly positive AUC difference (0.05 and 0.08, *p* < 0.001) and significantly positive SD difference (2.24 and 2.06, *p* < 0.001). *ω*_2_ = 0 and 0.1 only provided AUC higher than 0.8 in the case of *Se* = 3, whereas *ω*_2_ = 0.7 and 0.9 failed in all cases and even the median values of Dmin were significantly larger (median values around 2-3 cm) than other cases. Based on these results, we decided to consider only the depth weighting values *ω*_2_ = 0.3 and 0.5 for depth weighting MNE in the comparisons with with MEM reconstructions.

**Fig. S1.**
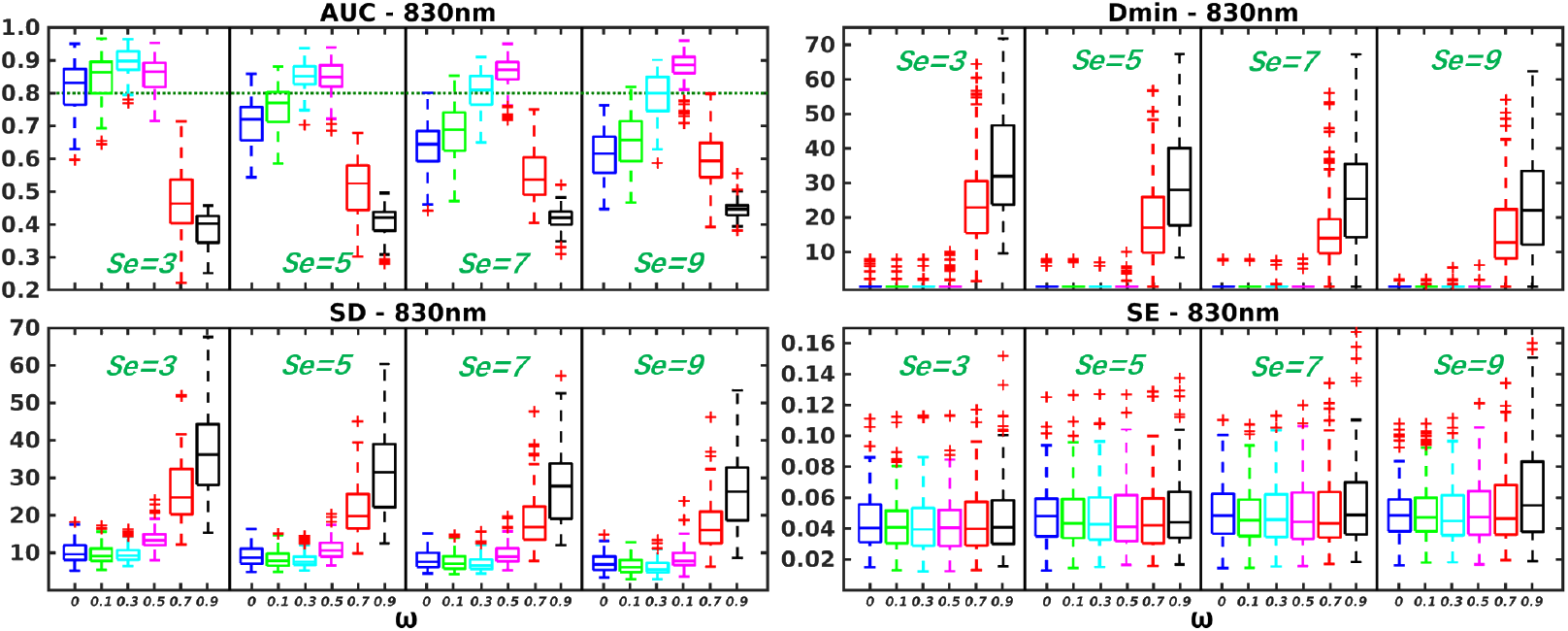
Evaluation of the performances of depth weighted MNE for different depth weighting factors *ω* = 0, 0.1, 0.3, 0.5,0.7, 0.9. Distribution of validation metrics (AUC, Dmin, SD and SE) are displayed using boxplot representations, for simulations involving superficial seeds only and for spatial extents *Se* = 3, 5, 7, 9.

### S3. MEM v.s. MNE with realistic simulations involving middle and deep seeds

In Fig. S2 and Table. S1, we are presenting the comparison of MEM and MNE in middle seeds case. First of all, we found that more depth compensation was required to provide good reconstructions in all scenarios. Thus, MEM(0.5,0.5) was compared to the best of MNE - MNE(0.5). Non-significant AUC and Dmin differences were found between them. However, MEM(0.5,0.5) provided significant lower SD than MNE(0.5), median value of difference of *SD* = −5.33, −4.80, −5.00, −4.95, *p* < 0.001 for *Se* = 3, 5, 7, 9 respectively. Fig. S3 and Table. S2 are presenting the comparison of MEM and MNE in the comparison of them in deep seeds case. Similarly, no significant AUC and Dmin differences were found. MEM(0.5,0.5) provided significant lower SD than MNE(0.5), median value of difference of *SD* = −6.39, −6.33, −6.97, −5.52, *P* < 0.001 for *Se* = 3, 5, 7, 9 respectively. For temporal performance in these two cases, similar to Fig. 3, MNE(0.5) gave significant lower SE (−0.01 or −0.02, *p* < 0.001) than MEM when *Se* = 3, 5 (small difference). No significant different SE was found in *Se* = 7, 9.

**Fig. S2.**
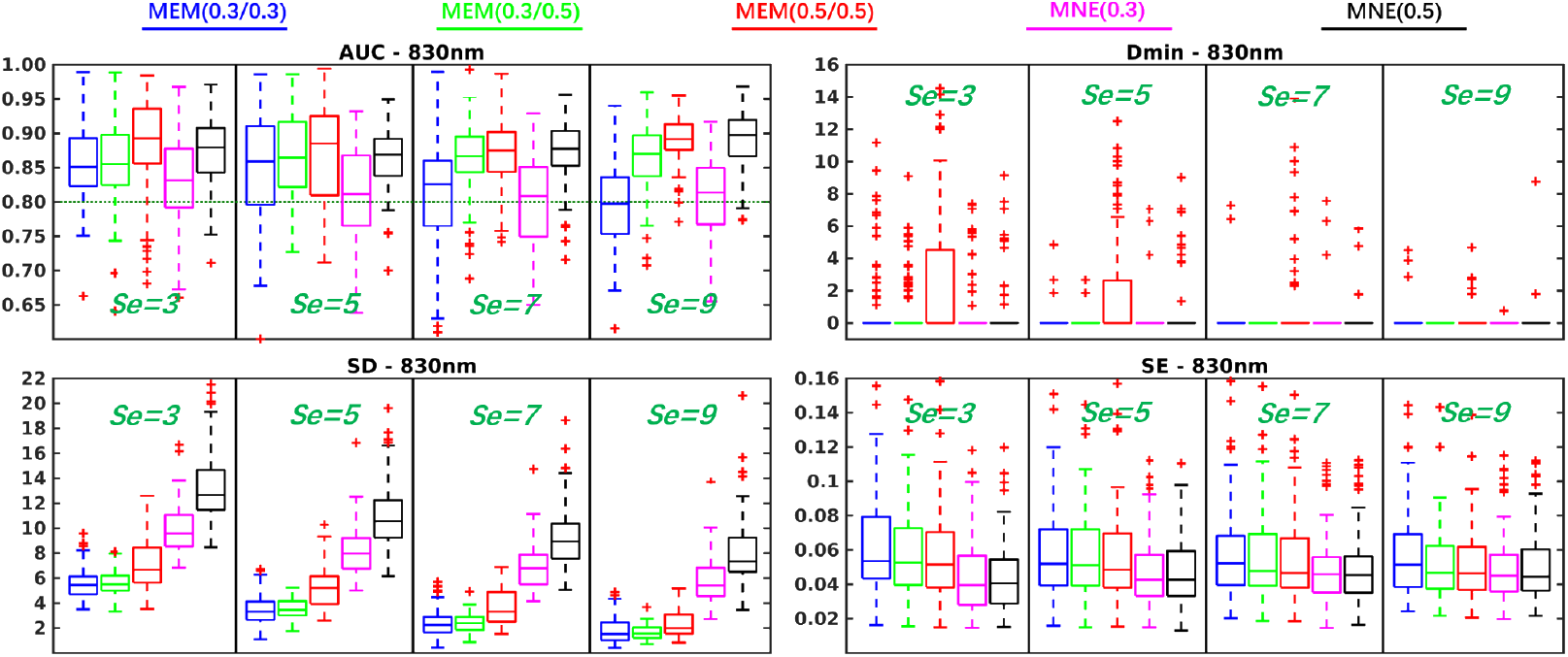
Evaluation of the performances of MEM and MNE using realistic simulations involving middle seeds for different spatial extent (*Se* = 3, 5, 7, 9). Boxplot representation of the distribution of four validation metrics for three depth weighted strategies of MEM and two depth weighted strategies of MNE, namely: MEM(0.3, 0.3) in blue, MEM(0.3, 0.5) in green, MEM(0.5, 0.5) in red, MNE(0.3) in magenta and MNE(0.5) in black. Results were obtained after DOT reconstruction of 830*nm* Δ*OD*.

**Table. S1.** Wilcoxon signed rank test results of reconstruction performance comparison of MEM and MNE in middle seeds case. Median values of paired difference are presented in the table. *p* values were corrected for multiple comparisons using Bonferroni correction, * indicates *p* < 0.01 and ** represents *p* < 0.001. Median of the paired difference of each validation metrics is color coded as follows: green: MEM is significantly better than MNE, red: MNE is significantly better than MEM and gray: non-significance.

| Middle Seeds |  | Se = 3 |  | Se = 5 |  | Se = 7 |  | Se = 9 |  |
| --- | --- | --- | --- | --- | --- | --- | --- | --- | --- |
|  |  | *MNE (0.3)* | *MNE (0.5)* | *MNE (0.3)* | *MNE (0.5)* | *MNE (0.3)* | *MNE (0.5)* | *MNE (0.3)* | *MNE (0.5)* |
| AUC | *MEM (0.3, 0.3)* | 0.03** | -0.03 | 0.03** | 0.00 | 0.02 | -0.05** | -0.02 | -0.10** |
|  | *MEM (0.3, 0.5)* | 0.03** | -0.03 | 0.05** | 0.01 | 0.05** | -0.01 | 0.05** | -0.02** |
|  | *MEM (0.5, 0.5)* | 0.06** | 0.02 | 0.07** | 0.01 | 0.07** | -0.01 | 0.08** | 0.00 |
| Dmin | *MEM (0.3, 0.3)* | 0.00 | 0.00 | 0.00 | 0.00 | 0.00 | 0.00 | 0.00 | 0.00 |
|  | *MEM (0.3, 0.5)* | 0.00 | 0.00 | 0.00 | 0.00 | 0.00 | 0.00 | 0.00 | 0.00 |
|  | *MEM (0.5, 0.5)* | 0.00 | 0.00 | 0.00 | 0.00 | 0.00 | 0.00 | 0.00 | 0.00 |
| SD | *MEM (0.3, 0.3)* | -4.05** | -7.21** | -4.27** | -7.25** | -4.10** | -6.40** | -3.58** | -5.43** |
|  | *MEM (0.3, 0.5)* | -4.00** | -7.06** | -4.09** | -6.90** | -3.96** | -6.40** | -3.65** | -5.45** |
|  | *MEM (0.5, 0.5)* | -2.54** | -5.33** | -2.46** | -4.80** | -2.85** | -5.00** | -3.08** | -4.95** |
| SE | *MEM (0.3, 0.3)* | 0.02** | 0.02** | 0.01** | 0.01** | 0.01** | 0.01* | 0.00 | 0.01 |
|  | *MEM (0.3, 0.5)* | 0.01** | 0.01** | 0.01** | 0.01** | 0.00 | 0.00 | 0.00 | 0.00 |
|  | *MEM (0.5, 0.5)* | 0.01** | 0.01** | 0.01** | 0.01* | 0.00 | 0.00 | 0.00 | 0.00 |

**Fig. S3.**
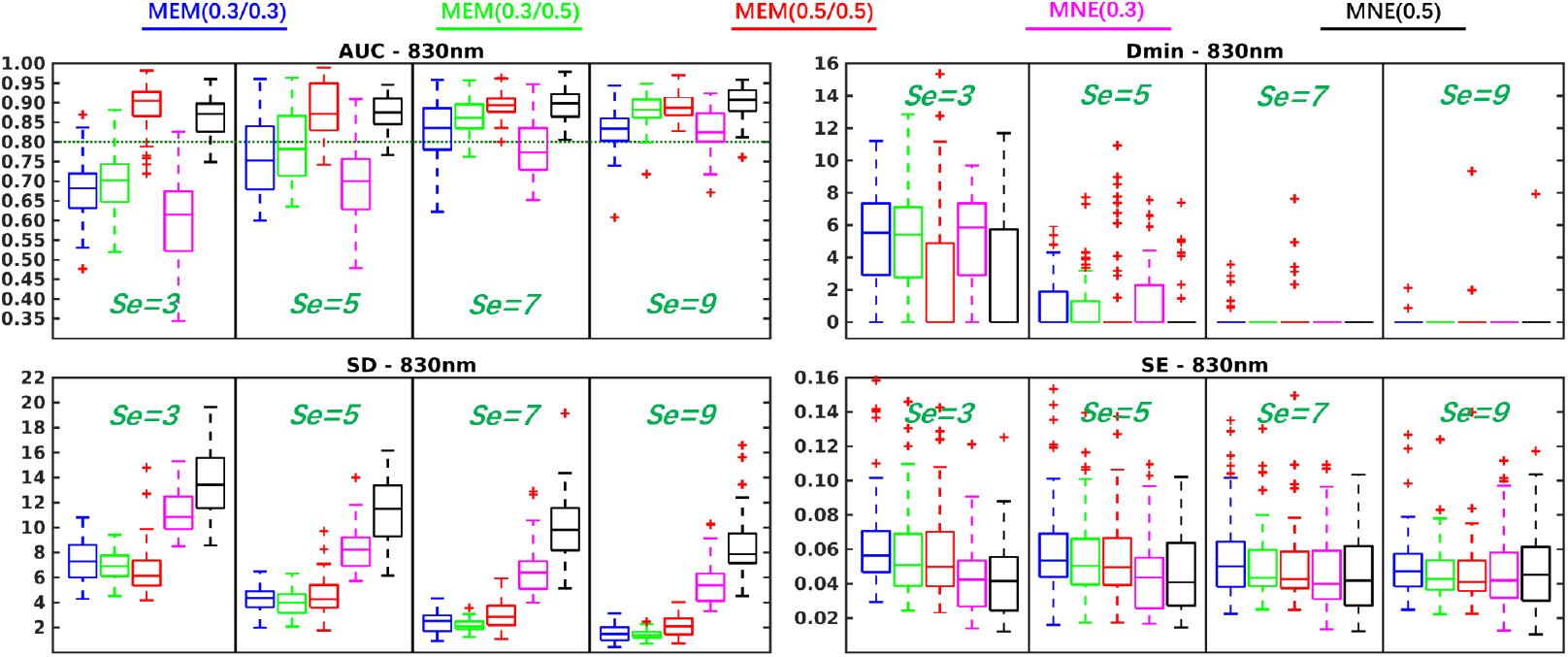
Evaluation of the performances of MEM and MNE using realistic simulations involving deep seeds for different spatial extent (*Se* = 3, 5, 7, 9). Boxplot representation of the distribution of four validation metrics for three depth weighted strategies of MEM and two depth weighted strategies of MNE, namely: MEM(0.3,0.3) in blue, MEM(0.3, 0.5) in green, MEM(0.5, 0.5) in red, MNE(0.3) in magenta and MNE(0.5) in black. Results were obtained after DOT reconstruction of 830*nm* Δ*OD*.

**Table. S2.**
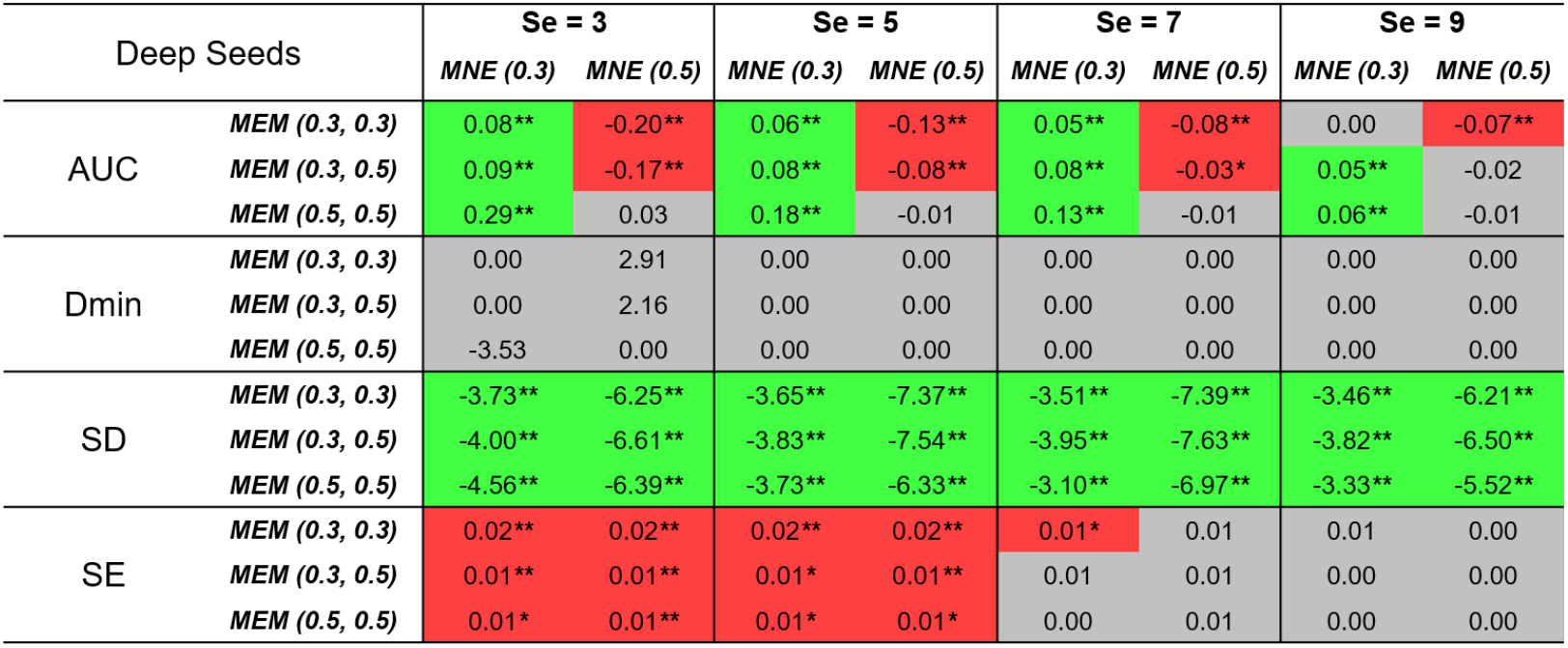
Wilcoxon signed rank test results of reconstruction performance comparison of MEM and MNE in deep seeds case. Median values of paired difference are presented in the table. *p* values were corrected for multiple comparisons using Bonferroni correction, * indicates *p* < 0.01 and ** represents *p* < 0.001. Median of the paired difference of each validation metrics is color coded as follows: green: MEM is significantly better than MNE, red: MNE is significantly better than MEM and gray: non-significance.

| Deep Seeds |  | Se = 3 |  | Se = 5 |  | Se = 7 |  | Se = 9 |  |
| --- | --- | --- | --- | --- | --- | --- | --- | --- | --- |
|  |  | *MNE (0.3)* | *MNE (0.5)* | *MNE (0.3)* | *MNE (0.5)* | *MNE (0.3)* | *MNE (0.5)* | *MNE (0.3)* | *MNE (0.5)* |
| AUC | *MEM (0.3, 0.3)* | 0.08** | -0.20** | 0.06** | -0.13** | 0.05** | -0.08** | 0.00 | -0.07** |
|  | *MEM (0.3, 0.5)* | 0.09** | -0.17** | 0.08** | -0.08** | 0.08** | -0.03* | 0.05** | -0.02 |
|  | *MEM (0.5, 0.5)* | 0.29** | 0.03 | 0.18** | -0.01 | 0.13** | -0.01 | 0.06** | -0.01 |
| Dmin | *MEM (0.3, 0.3)* | 0.00 | 2.91 | 0.00 | 0.00 | 0.00 | 0.00 | 0.00 | 0.00 |
|  | *MEM (0.3, 0.5)* | 0.00 | 2.16 | 0.00 | 0.00 | 0.00 | 0.00 | 0.00 | 0.00 |
|  | *MEM (0.5, 0.5)* | -3.53 | 0.00 | 0.00 | 0.00 | 0.00 | 0.00 | 0.00 | 0.00 |
| SD | *MEM (0.3, 0.3)* | -3.73** | -6.25** | -3.65** | -7.37** | -3.51** | -7.39** | -3.46** | -6.21** |
|  | *MEM (0.3, 0.5)* | -4.00** | -6.61** | -3.83** | -7.54** | -3.95** | -7.63** | -3.82** | -6.50** |
|  | *MEM (0.5, 0.5)* | -4.56** | -6.39** | -3.73** | -6.33** | -3.10** | -6.97** | -3.33** | -5.52** |
| SE | *MEM (0.3, 0.3)* | 0.02** | 0.02** | 0.02** | 0.02** | 0.01* | 0.01 | 0.01 | 0.00 |
|  | *MEM (0.3, 0.5)* | 0.01** | 0.01** | 0.01* | 0.01** | 0.01 | 0.01 | 0.00 | 0.00 |
|  | *MEM (0.5, 0.5)* | 0.01* | 0.01** | 0.01* | 0.01* | 0.00 | 0.01 | 0.00 | 0.00 |

